# Distinct geometries, comparable interfaces: binding modes and thermodynamic implications in conventional and single domain antibodies

**DOI:** 10.64898/2026.06.01.729307

**Authors:** Antoine Hauser, Gauthier Dangla Pélissier, Frédéric Cazals

## Abstract

Heavy-chain only antibodies, produced by the adaptive immune systems of camelids and cartilaginous fish, complement canonical antibodies comprising both heavy and light chain variable domains. Using an integrated interface model that unifies interfacial atoms–including solvent molecules, contacts, buried surface areas, and interface curvature measures, we shed light on two aspects of antibody binding that have so far remained elusive when comparing single domain (SdAb) and double domain (DdAb) antibodies.

First, contrary to previous reports of smaller SdAb interfaces, we show that SdAb achieve an interface size comparable to that of DdAb despite using a single variable domain and fewer interface residues, a consequence of a geometric pattern driven by convexity and curvature effects. Second, we show that SdAb exhibit a broader diversity of binding modes than previously reported, with a prominent role played by FR regions.

Finally, we discuss the thermodynamic implications of these findings for the design of high-affinity single domain antibodies, with particular relevance to protein engineering and design.

## 1 Introduction

### 1.1 Antibodies and their binding modes

#### Antibodies, structure and evolution

Antibodies (Abs) are central molecules of the adaptive immune system, recognizing pathogens and contributing to the immune defense. Abs come in different isotypes that differ in the number of subunits (polypeptide chains) and their domains. For example, the so-called IgG are heterotetramers involving two heavy and two light chains, resembling a Y-shaped molecule. Ab chains are structured into domains based on the underlying genes. While constant domains are encoded in constant (C) gene segments, variable domains are encoded in V(D)J gene segments, which give rise to somatic hypermutations to define Ab lineages. In Ab, the N-terminal domain of each chain is the variable domain. This variable domain itself can be divided into *framework* regions (FRs) and *complementarity determining* regions (CDRs). In canonical Ab, a variable domain comprises three CDRs interspersed with four FRs, in the order FR1-CDR1-FR2-CDR2-FR3-CDR3-FR4. Ab constitute a wide and complex class of molecules, and the evolution/classification of isotypes has been undertaken based on conservation properties of constant domains [1, 2, 3]. In this work, we compare the binding modes of two classes of Ab molecules: Double variable domain Ab (DdAb) and Single variable domain Ab (SdAb). The first, DdAb, also called *canonical Ab*, uses two variable domains (VD) denoted VH and VL respectively–located on the heavy and light chains respectively, to bind the antigen. The second, SdAb, uses a single variable domain. The VD of the latter is often called a nanobody.

#### Binding: the case of canonical Abs

The binding of an Ab to its antigen is a complex phenomenon. In general, it is qualified in terms of binding affinity – a thermodynamic quantity expressed say in kcal/mol, avidity – a quantity assessing multivalent binding, and specificity – ability to engage several partners with a sufficient affinity. Understanding binding remains challenging due to the complexity of Ab molecules and the various binding situations. Let us discuss it say for an IgG. The classical vision of Ab binding devotes a key role to the CDR3, and specific mutations may enhance affinity by pre-conditioning the binding geometry and lowering the so-called entropic penalty upon binding [4, 5]. Beyond CDR3, all CDRs usually play a significant role in defining the interface, with in general comparable contributions of the first two CDRs, CDR1 and CDR2, and the CDR3 [6]. Beyond variable domains, the role of constant domains has also been characterized by comparing Ab with identical variable domains yet different constant domains [7]. Hinge flexibility and antibody geometry may also condition the ability to reach cluttered regions–*e.g.* broadly neutralizing Ab targeting the stem of spike proteins in viruses [8]. Constant domains also influence the orientation of binding and the spacing between Fab arms, which influences avidity in case of multivalent binding [9, 10]. Finally, glycosylation, a post-translational modification consisting of attaching carbohydrate chains (glycans) to specific amino acids (mainly in the Fc region) is also known to play a crucial role in the immune function [11].

#### From canonical Abs to VHH

The adaptive immune systems of camelids and cartilaginous fish generate homodimeric Abs consisting of heavy chains only, referred to as heavy chain only antibodies (Fig. S1). Despite the absence of clear evidence supporting an evolutionary preference for either antibody type, the genetic divergence that gave rise to structurally distinct antibody formats, coupled with their functional convergence in distantly related lineages, suggests that comparable selective pressures have shaped their evolution.

Four conditions to obtain an effective homodimeric H-chain antibody have been identified [1]: (i) absence of CH1 domain to prevent retention in the endoplasmic reticulum; (ii) ability to generate an extensive and diverse primary repertoire; (iii) solubility in the absence of a light-chain variable region; (iv) possible stabilization via disulfide bonds. From a functional standpoint, it has also been speculated that smaller, prolate-shaped Ab could complement canonical ones whose interface tend to be flat or concave [1]. An exhaustive comparison of SdAb and DdAb antibodies and their binding modes has been carried out in various recent studies [12, 13, 14, 15]. These works have focused on various parameters including the length and composition of FR and CDR regions, the structural diversity of CDRs, the characterization of epitopes, paratopes, the binding modes, and the amino acids composition biases. We summarize them below.

These studies are especially important since application-wise, single domain Abs have a growing repertoire of uses both in clinical and basic research, a testament to their complement of the conventional VH/VL antibodies. As of 2025, about 200 Ab-based therapeutics have been approved by different health authorities to treat various pathologies [16].

#### Binding: thermodynamics and kinetics

From a physical chemistry standpoint, the non covalent interaction of two molecules is qualified by the association and dissociation free energies [17], namely Δ*G_d_* = −*RT* log *K_d_/c*^0^ and Δ*G_a_* = −*RT* log *c*^0^*K_a_* = *RT* log *K_d_/c*^0^ = Δ*H* − *T* Δ*S*. Thus, by nature, the affinity has an enthalpic component (Δ*H*) qualifying the interaction energy, but also an entropic component (*T* Δ*S*) qualifying the loss of dynamical properties upon complex formation. The formation of the complex indeed restricts the degrees of freedom of both partners. These two competing interests illustrate the enthalpy - entropy compensation phenomenon [18, 19], which stipulates that a favorable enthalpic change upon association is accompanied by an entropic penalty. Thereafter, a special care is devoted to link the structural parameters analyzed to thermodynamics and kinetics.

### 1.2 SdAb versus DdAb: Established features

The following findings have been reported recently [12, 13, 14, 15].

#### FR and CDR sequences: conservation and length

Framework (FR) regions tend to be more conserved in SdAb [13]. Using the normalized mean Hamming distance to compare sequence diversity, CDR-H1 are equally diverse in SdAb and DdAb, while CDR-H2 (resp. CDR-H3) are less (resp. more) diverse in SdAb than in DdAb [13]. While lengths of FR regions are essentially conserved, CDR-H3 is longer by ∼ 3-4 residues in SdAb [13]. Despite this difference in length, the overall amino acid composition of the CDRs remains broadly comparable between the two antibody types although differences occur at individual positions [12, 15].

#### Structural diversity of CDR regions

Loop geometry in SdAb vs DdAb has been studied by computing backbone lRMSD for CDR-H1, CDR-H2 and CDR-H3. It appears that SdAb loops are structurally more diverse than their sequence diversity alone predicts: H1: ∼ 3.36Å vs 1.60 Å ; *H*2: 1.86 Å vs 1.78Å; H3: 6.43Å vs 3.52Å [13]. Without VL hindrance, longer CDR-H3 in SdAb explore more diverse geometries and orientations, creating new binding opportunities [12, 14, 15]. This picture can be refined by also considering a fourth CDR-like loop, sometimes referred to as CDR4 [20]. This loop joins the *β* strand D and E and lies spatially beside CDR1 and CDR2, with which it interacts.

#### Paratopes and epitopes

Previous work has reported a BSA of 768 ± 201Å^2^ for SdAb and 833 ± 259Å^2^ for DdAb (VH+VL) [14]. It has also been reported that interfaces of SdAb:antigen and DdAb:antigen use a *consensus paratope* of roughly comparable size, namely ∼ 57 positions for VH+VL in DdAb and ∼ 50 positions for the VD of SdAb [13]. A smaller size for SdAb paratopes was also observed in [15], using two paratope definitions (distance based threshold of 4.5Å: 3.6 residues less; and interaction based definition: 2.6 residues less). These results provide a useful reference point for our analysis, but as explained in Sec. 2.3, suffer from the distance bias which prevents reporting selected atoms in contact even if they lose solvent accessibility. We recompute these quantities on our datasets with and without water molecules (Table 1) and reach different conclusions. On the antigen side, Gordon et al. reported modest differences: DdAb epitopes are slightly more linear, whereas SdAb epitopes are marginally less solvent-accessible according to their SASA-based accessibility metric, with substantial overlap between the distributions [15].

#### Interface packing, geometry, and binding modes

The residue-level contact density of an interface is defined as the mean number of partner residues contacted by each interfacial antibody residue. It has previously been reported that the contact density of SdAb residues is approximately 1.7 times greater than that of the combined VH and VL residues of DdAb, and approximately 1.4 times greater than that of DdAb VH residues alone [14].

Previous work has investigated interface geometry using the Lawrence–Colman shape complementarity index [21, 13, 14]. Using Connolly’s *G*^1^ surface, this index quantifies how closely the two interacting surfaces match by comparing the normals at paired surface points, with an exponential weighting based on their separation. Mitchell and Colwell reported a slightly higher mean shape complementarity for SdAb than for DdAb interfaces (0.72 versus 0.70), although the magnitude of the difference was small [14].

Using the molecular *i.e.* Connolly surface, the *topography i.e.* concavity/convexity has also been studied [22] using a fractal index defined as the slope of the linear function *log(number of atoms within a sphere centered on a surface point)* as a function of *log(sphere radius)* [23]. These studies and the subsequent ones have shown that SdAb paratopes can adopt convex geometries and insert a CDR into clefts, grooves, or pockets. However, these examples did not establish whether such geometries constitute a general SdAb binding mode [12]. Consistently, Gordon et al. found that SdAb epitopes were only marginally less solvent-accessible than DdAb epitopes, with substantial overlap between the two distributions [15].

Finally, binding modes characterized by the regions contacting the antigen have been studied [14], with CDR-H1+H2+H3 together accounting for ∼ 53% (resp. 16%) in DdAb (resp. SdAb) [14]).

The previous analyses may be complemented by observing interfacial residues do not display any clear statistical bias in terms of side chain size in SdAb (Fig. 1(Second row), Table S4), or biases in a.a. composition (Fig. S8; Tables S3). This suggests that functional diversity arises primarily from geometric and length variations rather than major shifts in residue physico-chemical properties.

**Figure 1:**
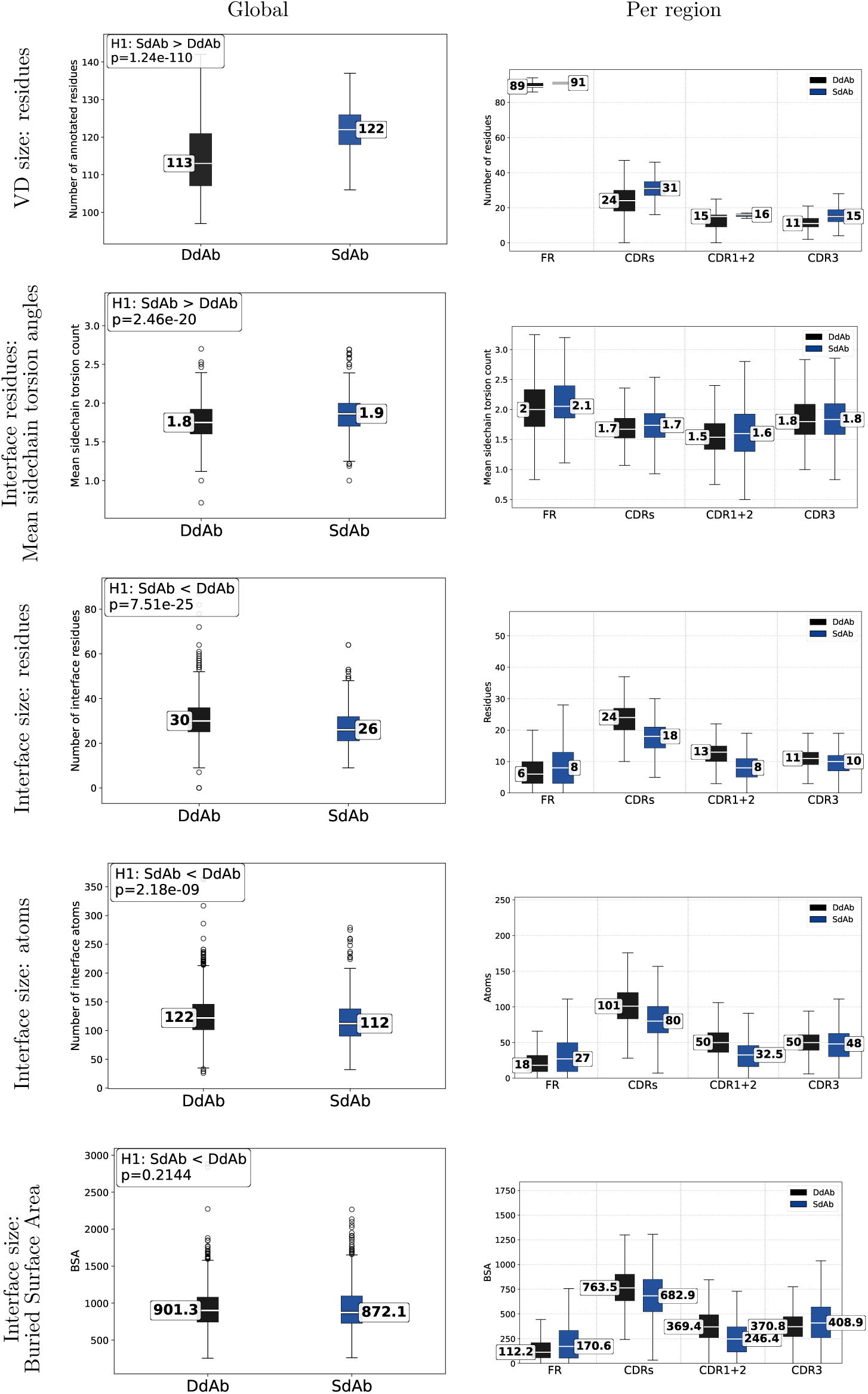
Dataset Ab-ALL, solvated models: comparison of SdAb vs DdAb interface properties. For DdAb VD size (first row), VH and VL are counted separately – one FAB casts two values. For the remaining rows, each FAB casts a single value. Mann-Whitney U tests were performed using one-sided and two-sided alternatives; the lowest p-value is displayed. SdAb has a smaller interface size than DdAb in term of residue or atom count, but SdAb shows a similar BSA, mainly from FRs and CDR3.

### 1.3 Contributions

The interfaces and binding modes of SdAb and DdAb have been studied in great detail, using various statistics and tools. We perform novel analyses using a model unifying interfacial atoms, contacts, buried surface areas, and curvature measures. Using this model, we show that both types of antibodies yield interfaces with distinct geometries, yet comparable in terms of buried surface area, a consequence of a geometric pattern driven by convexity and curvature effects. We conclude by discussing the thermodynamic and kinetic implications of these findings, which are critical issues in protein engineering and design.

## 2 Material and methods

### 2.1 Datasets

#### Structural databases, SdAb and DdAb

We use two databases for SdAb and DdAb (Fig. S1, Table S2). The first is IMGT/3Dstructure-DB, which provides curated antibody structures with standardized IMGT annotations ([24] and IMGT/3Dstructure-DB). The second is SAbDab-nano, a database dedicated to single-domain antibodies, including VHH and VNAR structures ([25]. (SAbDab-nano is a subset of SAbDab [26].)

#### Dataset construction and resolution stratification

Structural analyses were performed on curated datasets selected according to structure resolution (Fig. S2). The main structural dataset (Ab-ALL; 2051 structures) contains complexes solved at a resolution of 4Å or better, whereas the high-resolution dataset used for solvation analyses (Ab-HR; 320 structures) was curated using the same criteria with a resolution cutoff of 2Å. Structures were excluded according to the following criteria (details in Section S1.1.1): (i) complexes for which the combined length of all target chains was below 50 residues, to exclude short peptide targets that could introduce geometric biases; (ii) structures in which an SdAb was used as a crystallization chaperone; (iii) structures in which the heavy- and light-chain variable domains of an DdAb were connected by a linker; (iv) manually identified outliers; (v) PDB entries containing both SdAb and DdAb, which were excluded from comparative analyses. (vi) redundant complexes were filtered using both sequence and interface similarity, while sequence-similar antibody–target complexes displaying distinct interfaces were retained.

On the Ab-ALL dataset, this selection procedure identifies 2134 antibody–antigen interfaces (1328 DdAb and 806 SdAb), computed from 2051 antibody–antigen complexes (1271 DdAb and 780 SdAb).

#### Antibody identification and classification

Within Ab-ALL, we identify structures corresponding to antibodies (Sec. S1). Among these, 1271 correspond to double-domain antibodies (DdAb), involving both heavy- and light-chain variable domains, and 780 correspond to single-domain antibodies (SdAb). The species distribution differs markedly between the two classes (Fig. S3). DdAb are mainly derived from human (842 structures) and mouse (376 structures), reflecting the strong representation of canonical antibodies from commonly studied organisms in structural databases. In contrast, SdAb are overwhelmingly derived from camelid species (654 structures), consistent with their biological origin and the prominence of VHH antibodies in this clade.

### 2.2 Antigens observed in datasets

#### Diversity characterization

To compare the target landscapes associated with different antibody formats, we computed pairwise distances between targets involved in antibody:target complexes. Each target was represented by its amino acid chain sequences. For targets composed of several chains, all pairwise chain:chain distances were first computed using sequence alignment with a BLOSUM62 substitution matrix and gap penalties. Chains from the two targets were then optimally matched using a one-to-one assignment minimizing the total chain:chain distance using the Hungarian algorithm. This produced a target:target distance matrix.

Hierarchical clustering was then performed from this distance matrix, and the resulting dendrogram was cut at a fixed distance threshold (0.6). This threshold-based strategy was used instead of forcing a fixed number of clusters, because many targets are highly dissimilar and the pairwise distance distribution is strongly concentrated near large values.

The same distance matrix was also projected in two dimensions using classical multidimensional scaling (MDS) [27] (Fig. S4). The MDS coordinates are therefore used only as a visualization of the global distance structure, whereas the colors correspond to clusters obtained from the dendrogram. The percentages indicated on the MDS axes correspond to the fraction of positive classical MDS inertia explained by each axis, and should be used to assess how much of the distance structure is captured by the two-dimensional projection. (Also note that the 2D projection can create overlaps between clusters well separated in higher dimension.)

#### Observed target families

Cutting the dendrogram at a target-sequence distance threshold of 0.6 revealed several biologically coherent target families (Table S1). In the combined antibody dataset, the largest displayed clusters corresponded to recurrent viral glycoproteins, including coronavirus spike and HIV gp120/CD4 complexes, as well as classical or recurrent structural biology targets such as lysozyme, influenza hemagglutinin, GPCR-related complexes, GABA-A receptor complexes, and potassium channels. When the analysis was stratified by antibody format, some families were recovered in both subsets (Fig. S4B,C), most notably coronavirus spike and lysozyme, indicating that these targets are broadly represented across antibody formats. Other clusters showed a stronger format-associated pattern. HIV gp120/CD4 complexes, influenza surface glycoproteins, KcsA, and integrin *α*IIb/*β*3 were mainly observed among DdAb structures (Fig. S4B), whereas GPCR-related complexes, proteasome-associated assemblies, ricin, and GFP were more prominent among SdAb structures (Fig. S4C). This pattern is consistent with the frequent use of single-domain antibodies as crystallization or conformational stabilizers for membrane proteins and large molecular assemblies, while conventional antibodies dominate several viral antigen datasets.

Importantly, these clusters should be interpreted as sequence-based target families rather than interaction-mode clusters. This analysis also reveals clear biases in the dataset composition, which should be kept in mind when comparing interface statistics, notably the buried surface area. To limit the influence of these biases, we applied the dataset filtering and redundancy-reduction procedure described above.

### 2.3 The Voronoi interface model

#### Interface models

The analysis of contacts in previous work uses a distance cutoff of 4.5Å or 5Å to declare that two atoms from the partners are in contact [14, 15]. We use instead the Voronoi based model [28, 29] used in previous studies [30, 6, 31, 32]. This model relies on the power diagram (weighted Voronoi diagram) of atomic balls in the solvent accessible model (Fig. S5, Fig. S6). That is, the atomic radii [33], which span the range [*r*_min_ = 1.4Å*, r*_max_ = 1.9Å] are expanded by the radius of a water probe (*r_w_* = 1.4Å). As documented in the aforementioned papers, the Voronoi based model is threshold-less and selects a single layer of interface atoms on each partner, accommodating interfacial water molecules. In particular, a direct contact between two atoms of the partners (denoted *A* and *B* by convention) is called an *AB* contact; while a contact mediated by a water molecule sandwiched between the partners is termed an *AW-BW* contact. These two types of contacts define the AB and AW-BW interfaces respectively, and their union defines the ABW interface. Thus, the model identifies atoms and their contacts. The Voronoi facets dual of pairs in contact define a polyhedral surface separating the partners, and its connected components identify binding patches. The polyhedral surface also comes with an intrinsic notion of curvature based on the angles between Voronoi facets, thereby avoiding any smooth approximation.

In comparing our results against those from [14], the following remarks apply:

1. The Buried Surface Area and the number of interfacial counts are consistent in our work since both are derived from the same Voronoi diagram. But they are not in [14] since the definition of the BSA and that of interfacial atoms are independent.
2. Defining interfaces from a distance threshold raises two issues. First, using atomic radii in the aforementioned range [*r*_min_*, r*_max_], the maximum distance between two atoms for them to be found in contact in the Voronoi/*α*-complex model is equal to 2(*r*_max_ + *r_w_*) = 6.6Å. This is markedly larger than the 5Å threshold from [14]. Second, defining the neighbors of an atom as those found in a sphere of a given radius yields a bias towards the interface core–as opposed to the interface rim [28].
3. The Voronoi models identifies a single layer of interface atoms regardless of the position in the interface–core versus rim. It inherently yields more interface atoms (Fig. S6). As we will see, the larger number of interface atoms yields more complex contact profiles.

#### Applications to complexes with SdAb and DdAb

Applying the Voronoi model to complexes requires distinguishing partners–called *A* and *B* as noted above, interfaces, and their connected components (Fig. S7):

- A *partner* is a set of **connected** polypeptide chains. Partners with several chains are used in two guises: a Fab is a partner with two chains; an (obligate) heterodimer is also a partner.

For DdAb, the main antibody-level interface generally combines VH and VL as a Fab interface. For SdAb, it corresponds to the single variable domain.

- The *(Voronoi) interface* between two partners is the aforementioned polyhedral surface extracted from the Voronoi model. As noted above, interfaces come into two guises: with and without crystallographic water molecules.
- A *binding patch* is a connected component of an interface.

The study of interfaces for canonical Abs and VHH yields various cases (Fig. S7). To single them out, we consider the number of connected components (c.c.) of antigen chains. (The number of c.c. for VD is always one.) We then compute the Voronoi interfaces between all VD chains and contacted antigen connected components. For example, in (V2), the two antigen chains are connected, whence a single interface, but consisting of two patches. In (V3), there are two disconnected antigen chains, whence two interfaces, each with one binding patch. Or in (F3), the two antigen chains are disconnected, yielding two interfaces, each with one patch. Selected complexes display several spatially disconnected binding patches (Fig. S12).

#### Software

We use three programs provided by the Structural Bioinformatics Library [34] to carry out our analysis: (i) The case study from Fig. S7 is sorted out using the graph of contacts between chains delivered by the Interface finder. (ii) Interfaces are modeled using Intervor [30, 28]. (iii) The BSA is evaluated using certified software Vorlume [35].

## 3 Results

Our integrated Voronoi based interface model is instrumental to sharpen and contrast the previous findings summarized in Sec. 1.2

### 3.1 SdAb and DdAb: similar contact areas while using fewer interface residues and atoms

#### SdAb and DdAb interfaces have similar buried surface area (BSA)

The buried surface area (BSA) at an interface is possibly the most important parameter as it is well known that BSA has an excellent correlation with the number of interface atoms which contribute on average ∼ 10Å^2^ [36, 37]. Solvation is known to also play a key role at interfaces in general, and Ab-Ag interfaces in particular [38], as water molecules exhibit a dual role. On the one hand, the desolvation of the protein surfaces coming in contact contributes to increasing the solvent entropy. On the other hand, water molecules squeezed at the interface are packing enhancers and contributors to the hydrogen bonding network. It has also been reported that the interfacial water molecules significantly increase the BSA and contribute to merging interface patches [30].

First, we confirm that the inclusion of water increases BSA significantly as shown by the following values (Table 1). With the dataset Ab-HR, the mean/median BSA increases from 794/793 to 1064/1036 Å^2^ for DdAb, and from 795/733 to 1038/967 Å^2^ for SdAb; focusing on Ab-ALL, the mean/median BSA increases from 835/819 to 927/903 Å^2^ for DdAb, and from 834/784 to 934/873 Å^2^ for SdAb. Using Ab-HR, we observe a strict proportionality between the number of interfacial water molecules and the BSA, both for DdAb and SdAb—with respectively 34.9 and 32.5 water molecules on average. As expected, the counts are much lower for Ab-ALL and CAM-ALL. For these reasons, we proceed with the water inclusive model and the dataset Ab-ALL.

At a glance, comparison of SdAb vs DdAb yields the following observations (Fig. 1): VD are larger in SdAb (Fig. 1(First row)) while using a.a. slightly larger as seen from the number of side chain torsion angles (Fig. 1(Second row)); interfaces in SdAb involve fewer residues (Fig. 1(Third row)) and fewer interface atoms (Fig. 1(Fourth row)). However, in total, interfaces of SdAb and DdAb have comparable BSA (Fig. 1(Fifth row)). We note that this fact owes to more significant contributions from CDR3 and framework regions. Finally, we also note that the overall physico-chemical composition of the interfaces is broadly comparable (Fig. S8,Fig. S11). Although specific amino acids are enriched or depleted at SdAb interfaces (Table S4). We also note that when the BSA buried on the VD and target sides are added, the values obtained for SdAb and DdAb become even more similar (Fig. S10), an observation we will get back to.

To explain the previous observations, we now study three related quantities: the BSA, the density of contacts across the interface and the interface curvature.

#### SdAb interfaces achieve a higher BSA thanks to a higher contact density

For each side of the interface, the atomic contact density is defined as the mean number of atoms across the interface contacted by each interfacial atom, including water molecules. (Recall that this information is encoded in our Voronoi model.) On the VD side, the atomic contact density of SdAb interfaces is higher than that of DdAb interfaces (Fig. 2(C)); the same trend is observed across regions at the residue level (Fig. S9). Thus, although fewer VD atoms participate in SdAb interfaces, each of these atoms interacts, on average, with a larger number of atoms across the interface. This larger number of neighbors has a direct implication in terms of BSA: for most regions, particularly FR2 and CDR3, SdAb atoms bury more surface area than DdAb atoms for a comparable number of interfacial atoms (Fig. 3). While these values are of the same order of magnitude as the ∼ 10Å^2^ buried per atom typically observed at protein–protein interfaces [36, 37], we observe significant differences: CDR3 atoms bury on average 7.56Å^2^ in DdAb versus 8.97Å^2^ in SdAb, and FR2 atoms 4.57Å^2^ versus 6.85Å^2^, respectively. Together, these results indicate that the comparable BSA of SdAb and DdAb interfaces does not result from a comparable number of interfacial atoms, but from a denser organization of contacts on the SdAb VD side.

**Figure 2:**
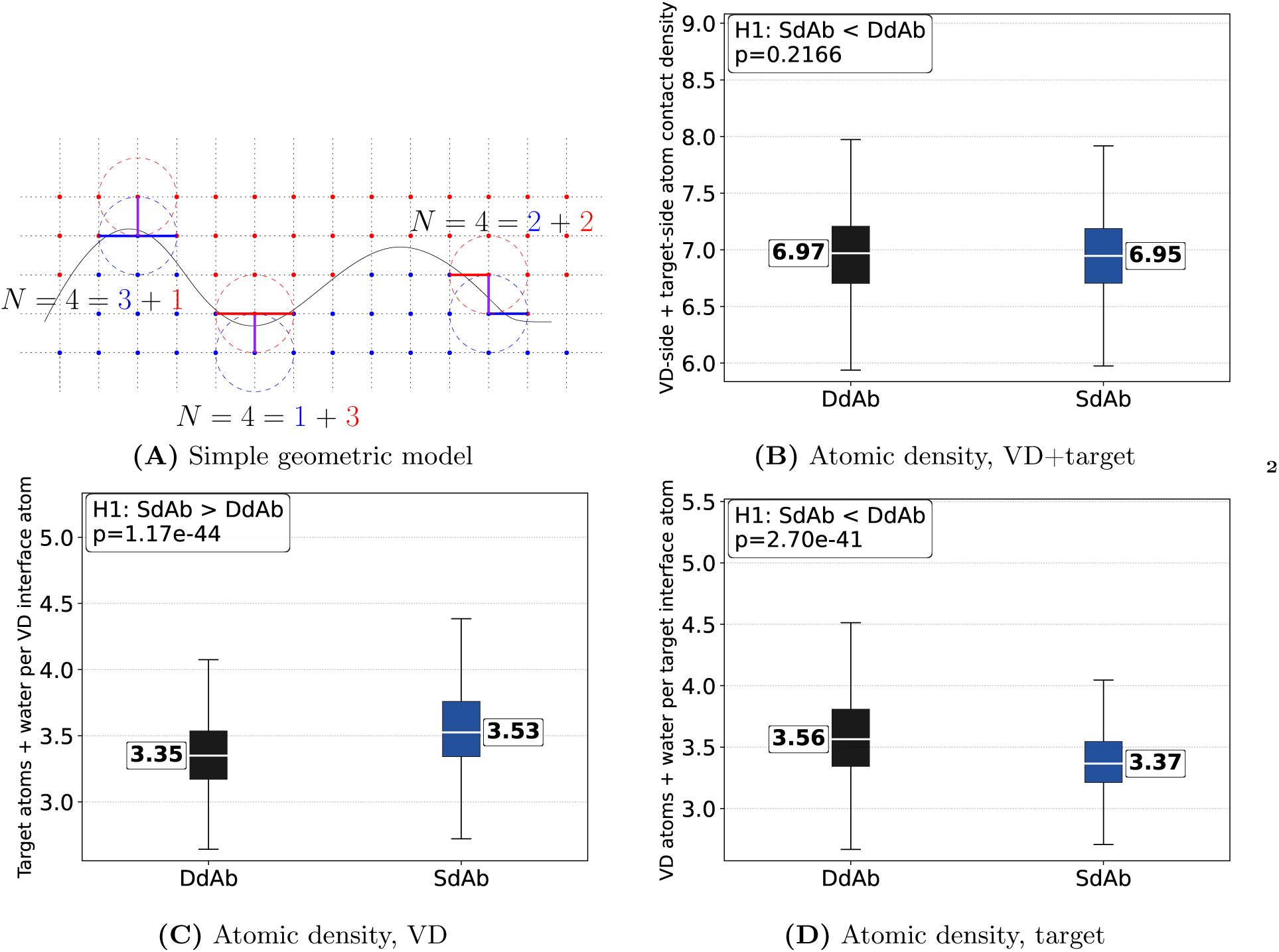
Dataset Ab-ALL, solvated models: atomic contact density at antibody–antigen interfaces. **(A)** Schematic explanation of the observed behavior: if atoms are similarly distributed locally on both sides of the interface, the summed atomic density is expected to remain approximately constant regardless of interface shape. **(B,C,D)** Atomic density defined as the average number of neighbors (partner+water molecules) per atom.

**Figure 3:**
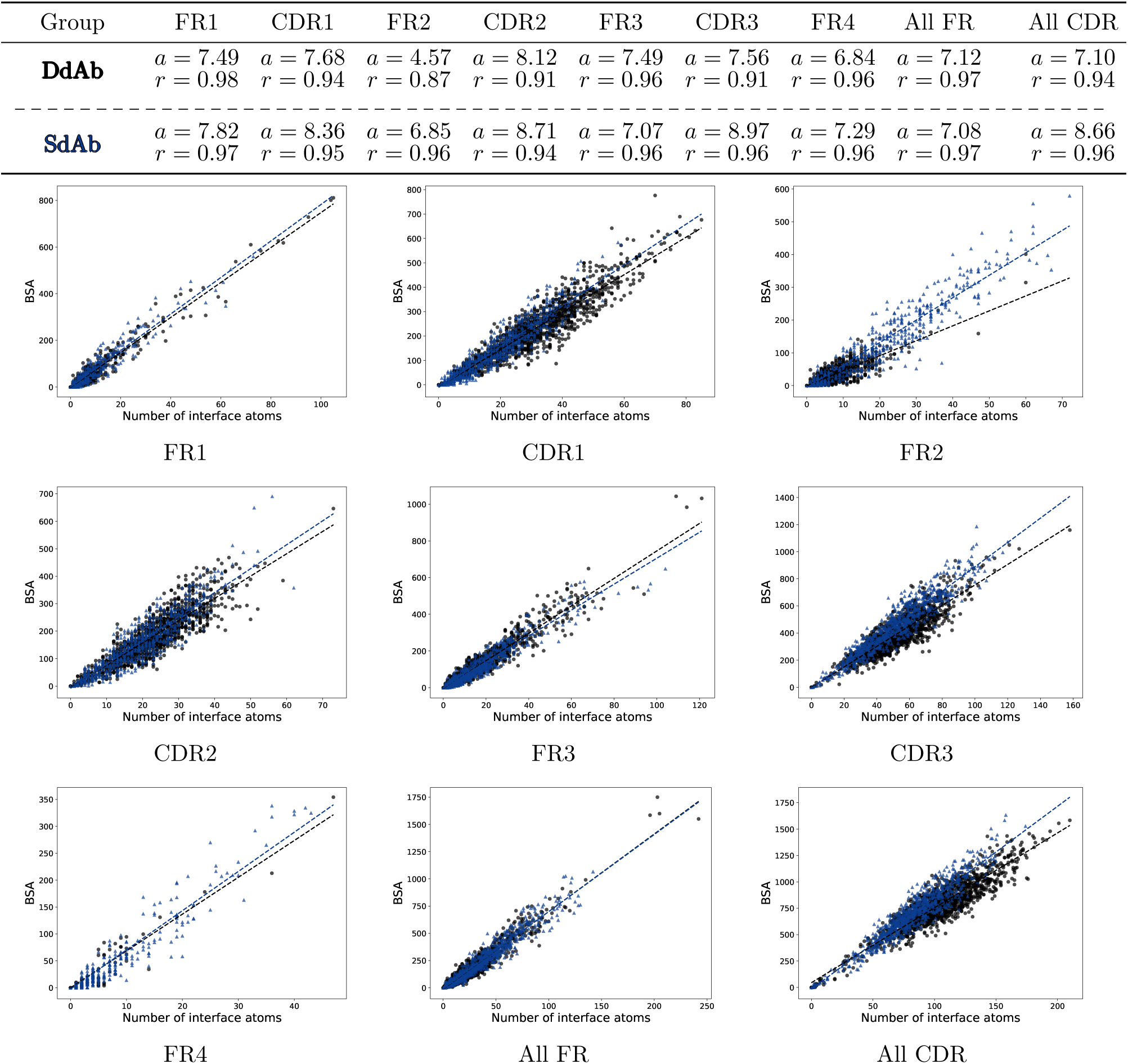
Relationship between buried surface area and the number of interfacial atoms across variable-domain regions. Scatter plots show the buried surface area (BSA) as a function of the number of atoms located at the interface for each framework and complementarity-determining region, using the Ab-ALL dataset and the solvated interface model. For most regions, particularly FR2 and CDR3, SdAb exhibit a larger variable-domain-side BSA for a given number of interfacial atoms than DdAb.

#### SdAb exhibit a higher contact density via convex interfaces

The previous facts turn out to be related to curvature effects and the pending question on whether SdAb form more convex interfaces [22].

Recall that the Voronoi interface model provides a notion of curvature based on angles between Voronoi facets (Fig. S5). The interface curvature can also be normalized by the interface area or the BSA. By convention, we observe an Ab:antigen interface from the antibody side, so that a positive (resp. negative) curvature indicates a convex (resp. concave) interface.

When comparing SdAb against DdAb interfaces, we note that both the signed curvature and the BSA-normalized signed curvature are shifted toward more positive values corresponding to convex paratopes (Fig. 4; significant differences were assessed by a positive one-sided Mann–Whitney *U* test). Geometrically, the convexity of the paratope exposes its atoms, which results in a higher contact density. We also note that while being statistically distinguishable, the curvature distributions overlap significantly with SdAb (resp. DdAb) exhibiting concave (resp. convex) interfaces (Fig. 4).

**Figure 4:**
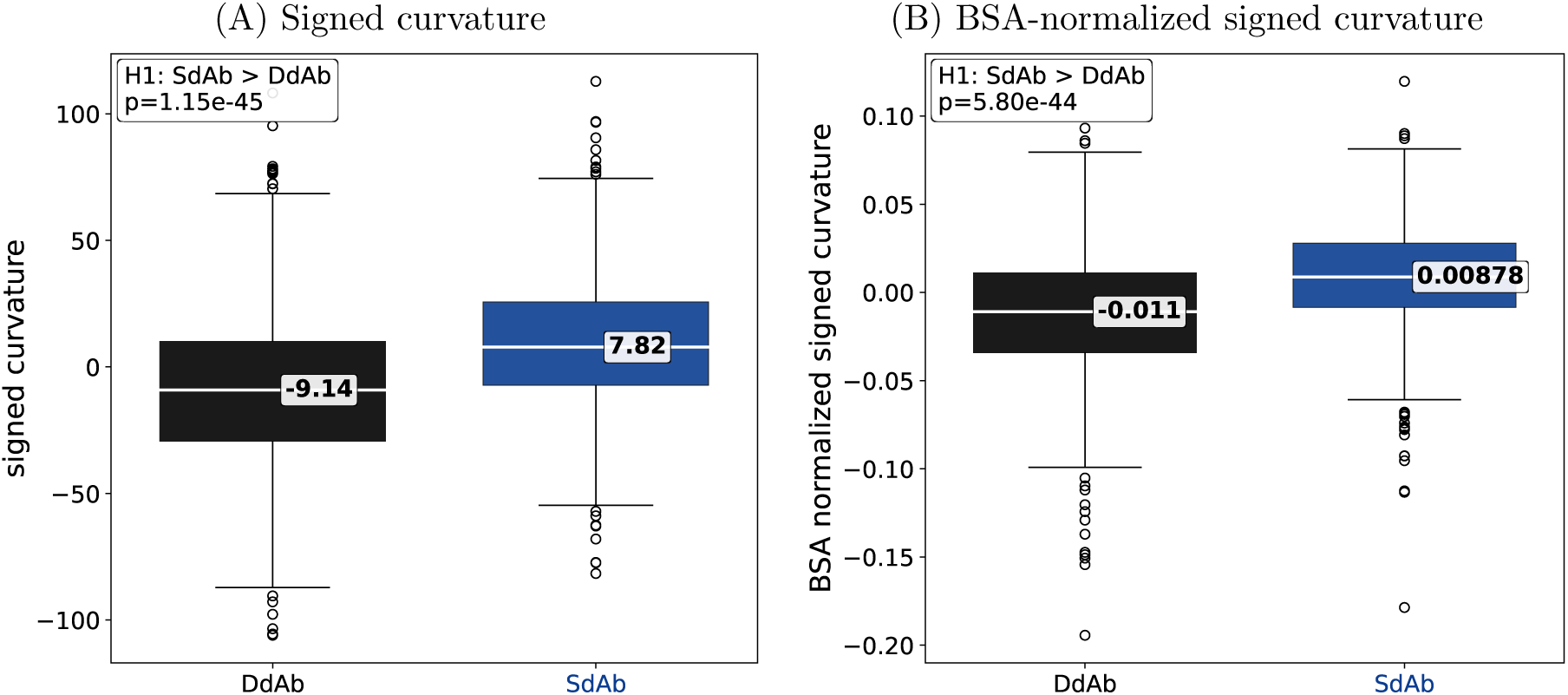
Comparison of antibody-side interface curvature between DdAb and SdAb. Dataset Ab-ALL, solvated model Positive signed curvature indicates a convex antibody-side interface, whereas negative curvature indicates a concave interface. (A) Signed curvature describes the overall interface geometry and may therefore depend on interface size. (B) Normalization by BSA measures signed curvature per unit of buried surface area. Both metrics are shifted toward more positive values for SdAb, showing that their binding surfaces are significantly more convex than those of DdAb. The persistence of the difference after BSA normalization shows that the greater convexity of SdAb interfaces is not solely explained by their larger buried surface area.

#### Closing the loop: BSA, contact density, and curvature effects

Finally, we establish a direct link between BSA and curvature by plotting the BSA against the number of interfacial atoms and the interfaces are stratified according to signed curvature (Fig. 5). Convex interfaces tend to bury more surface area per VD-side atom than concave interfaces, an effect which is particularly pronounced for CDR3–with coefficients equal to 8.66 (resp. 7.35) for convex (resp. concave) interfaces.

**Figure 5:**
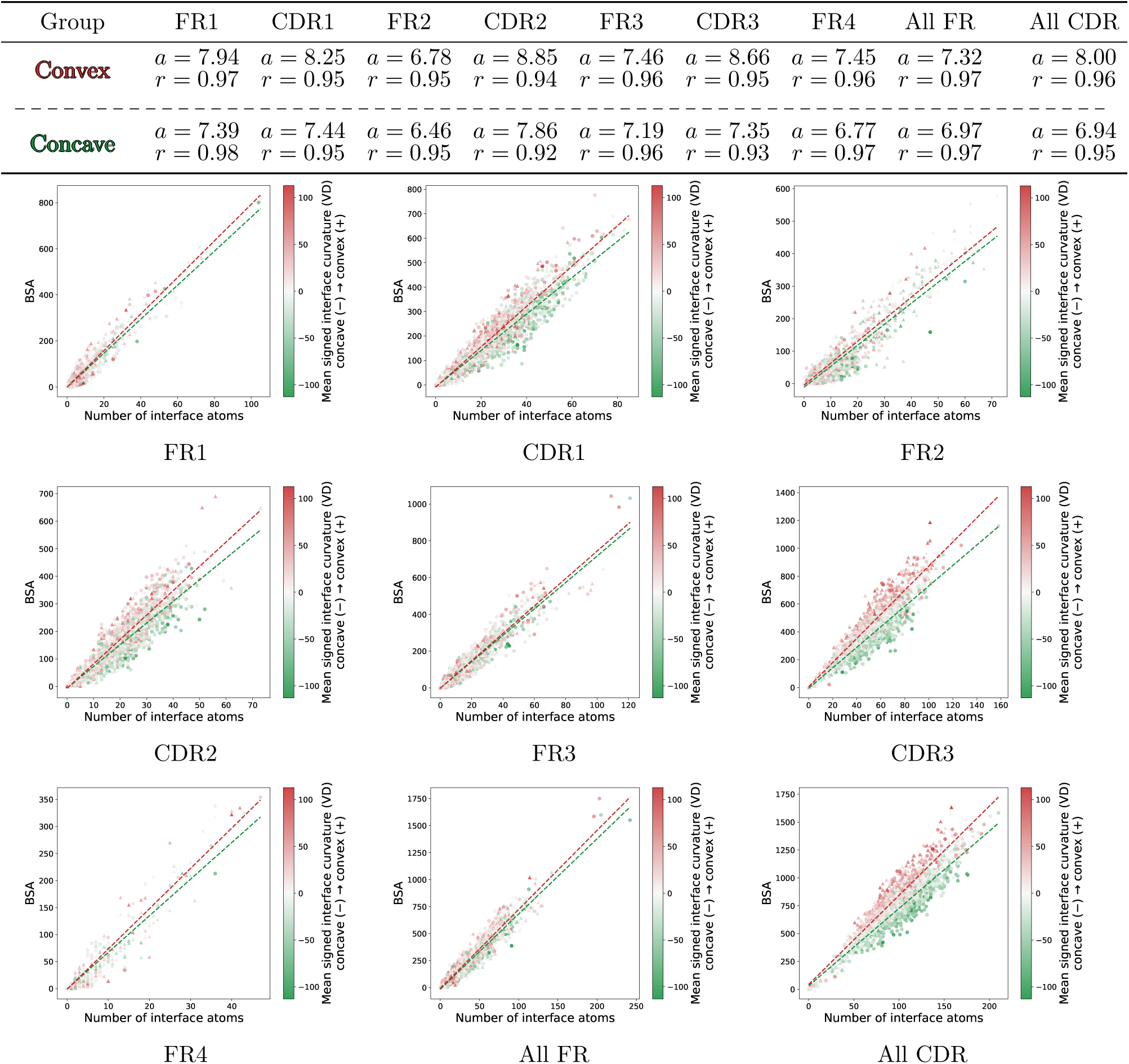
Relationship between buried surface area, interfacial atom count, and local interface curvature. Scatter plots show, for each region, the buried surface area (BSA) as a function of the number of interfacial atoms on the variable-domain side. Points are colored according to signed curvature, with convex interfaces (positive signed curvature) shown in red and concave interfaces (negative signed curvature) shown in blue. SdAb interfaces are represented as triangles and DdAb as disks. Separate linear regressions were fitted for convex and concave interfaces. Overall, atoms from convex interfaces tend to bury more surface area on the variable-domain side than atoms from concave interfaces, an effect that is particularly marked for CDR3.

We summarize these observations using a simple geometric model showing that the contact density is larger (resp. smaller) in convex (resp. concave) regions (Fig. 2(A)). On this model, atoms belonging to a protruding convex region contact a larger number of neighboring atoms than those belonging to a flat or concave region. The sum of atomic densities is however constant, a property which we observe in real complexes (Fig. 2(B)).

#### Biological illustrations

Despite the differences on interface organization (contact density, curvature), the two antibody formats achieve comparable total buried surface areas. To provide a structural perspective on these quantitative trends, we illustrate several representative and extreme binding geometries observed in our dataset (Fig. 6). In particular, these examples highlight the broad range of geometries accessible to SdAb and show that their ability to recognize clefts and pockets does not imply that they systematically target recessed or hidden epitopes. They include an SdAb with high positive curvature resulting from the deep insertion of a CDR into the antigen (Fig. 6(A)), an SdAb forming a concave lateral contact with its antigen (Fig. 6(B)), and an SdAb binding within a pocket that surrounds much of the variable domain (Fig. 6(C)). In contrast, the DdAb example in Fig. 6(D) illustrates a classical concave binding geometry in which the antigen is accommodated between the VH and VL domains.

**Figure 6:**
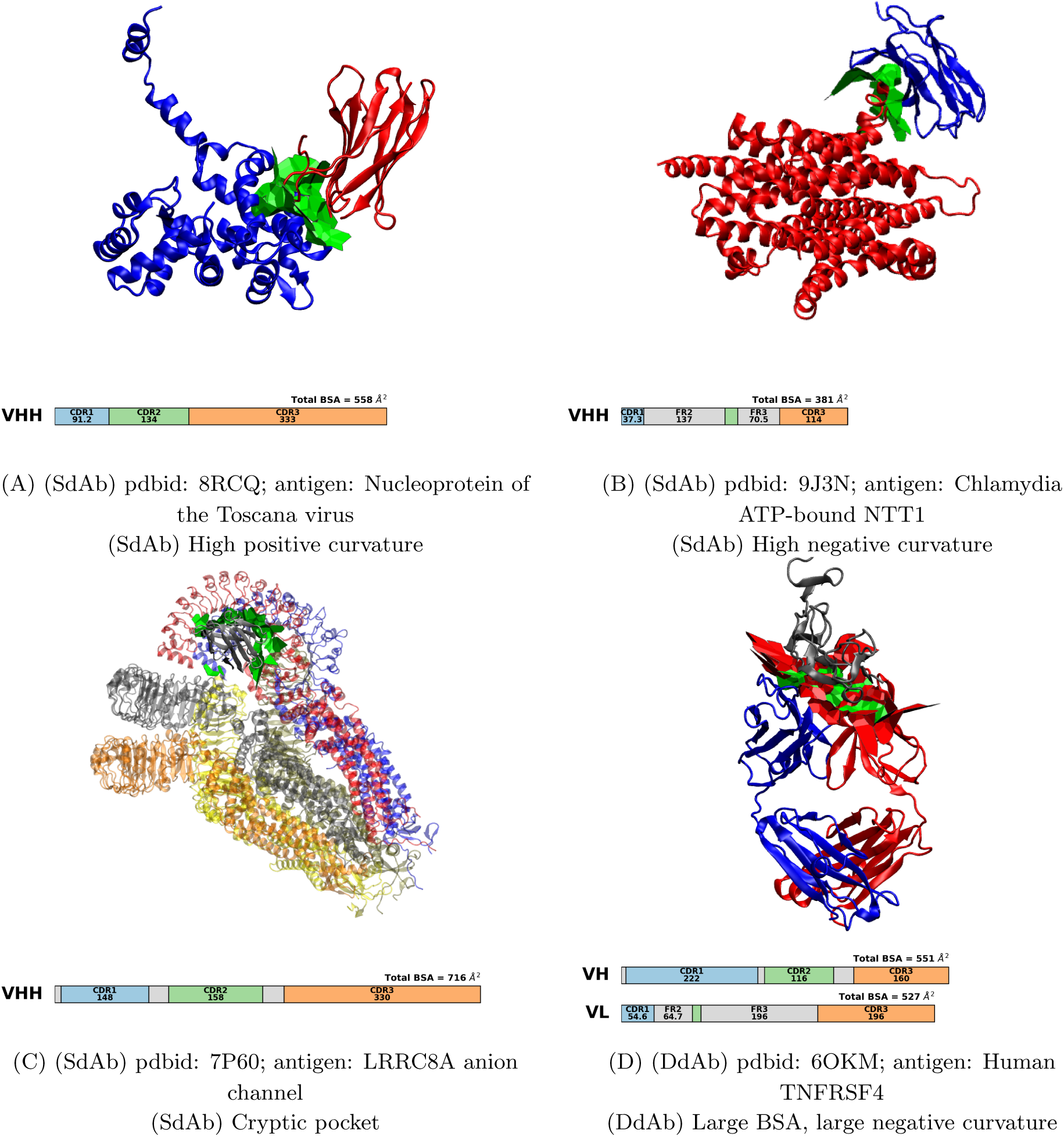
Extreme binding geometries curvature-wise, in SdAb and DdAb complexes. **(A) (B)** Two SdAb examples with extreme positive and negative curvature values, respectively. In (B), the interface also displays an unusual framework-mediated contact on the side of the variable domain, rather than being restricted to the canonical CDR face. **(C)** An SdAb binding a pocket of a large target, with contacts surrounding most of the variable domain. This binding geometry appears well suited to the compact architecture of single-domain antibodies, whereas a larger DdAb would likely be sterically constrained in accessing such a pocket. **(D)** An DdAb:antigen complex combining a large buried surface area and high curvature with an anisotropic interface involving the CDRs of both VH and VL chains.

The potential biological relevance of these distinct binding geometries is further illustrated by the prefusion RSV F glycoprotein. The 1G9 nanobody–a broadly neutralizing SdAb–binds a lateral, relatively membrane-proximal epitope at antigenic site IV [39], whereas the 32.4K Fab targets the membrane-distal apex at antigenic site ∅ [40] (Fig. 7). Thus, on the same antigen, the two antibody formats can exploit markedly different surface regions and binding geometries.

**Figure 7:**
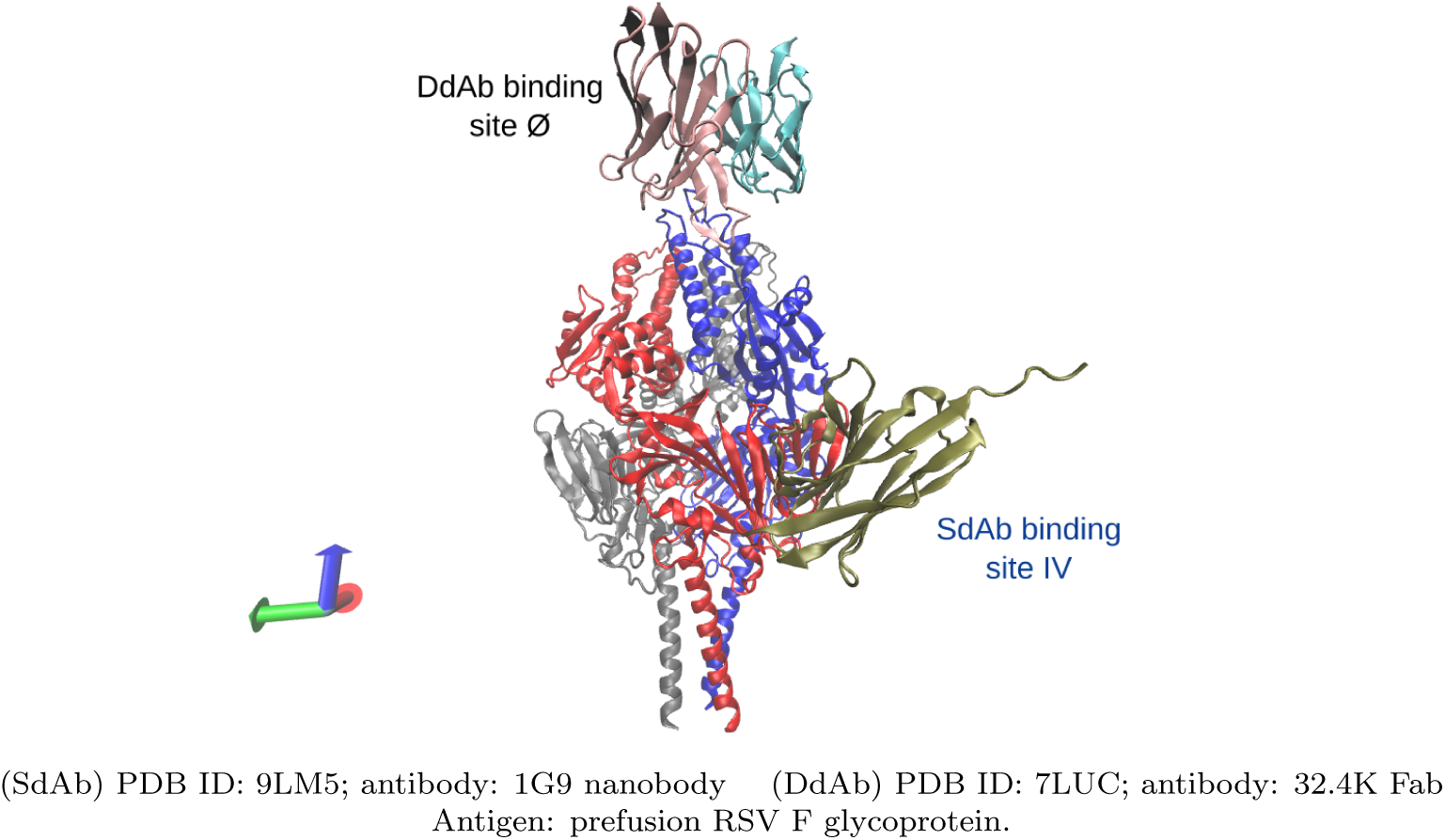
Example of distinct epitopes targeted by an SdAb and a DdAb on the same antigen. Superposition of SdAb–antigen and DdAb–antigen complexes involving the prefusion respiratory syncytial virus F glycoprotein. The SdAb 1G9 binds a lateral, relatively membrane-proximal epitope at antigenic site IV [39], whereas the DdAb 32.4K Fab binds the membrane-distal apex at antigenic site Ø [40]. This example illustrates that SdAbs can recognize relatively constrained epitopes, although this is not a systematic feature of SdAb binding.

### 3.2 SdAb vs DdAb: different binding modes with enhanced FR contributions

With three CDRs and four FRs, the number of possible subsets of regions of a VD at an interface is 2^7^ − 1 = 127. Call each subset a *profile*. Profiles are key to understand how the antibody interacts with its antigen. Previous work has counted 21 and 14 profiles for SdAb and DdAb, respectively [14]. In the following, we revisit and enrich the previously discussed binding modes, namely the canonical three-loop mode, the H3-dominated mode two-loop asymmetric modes (H1+H3 or H2+H3), as well as framework assisted modes.

#### Interface definitions explain discrepancies with previously reported contact profiles

As already explained, our Voronoi interface model, consistently with the definition of BSA, identifies more interface atoms (Sec. 2.3). Significant differences are observed in the contact profiles already published, and two of them deserve a special attention ([14, Fig. 3] vs Fig. 8)):

**Figure 8:**
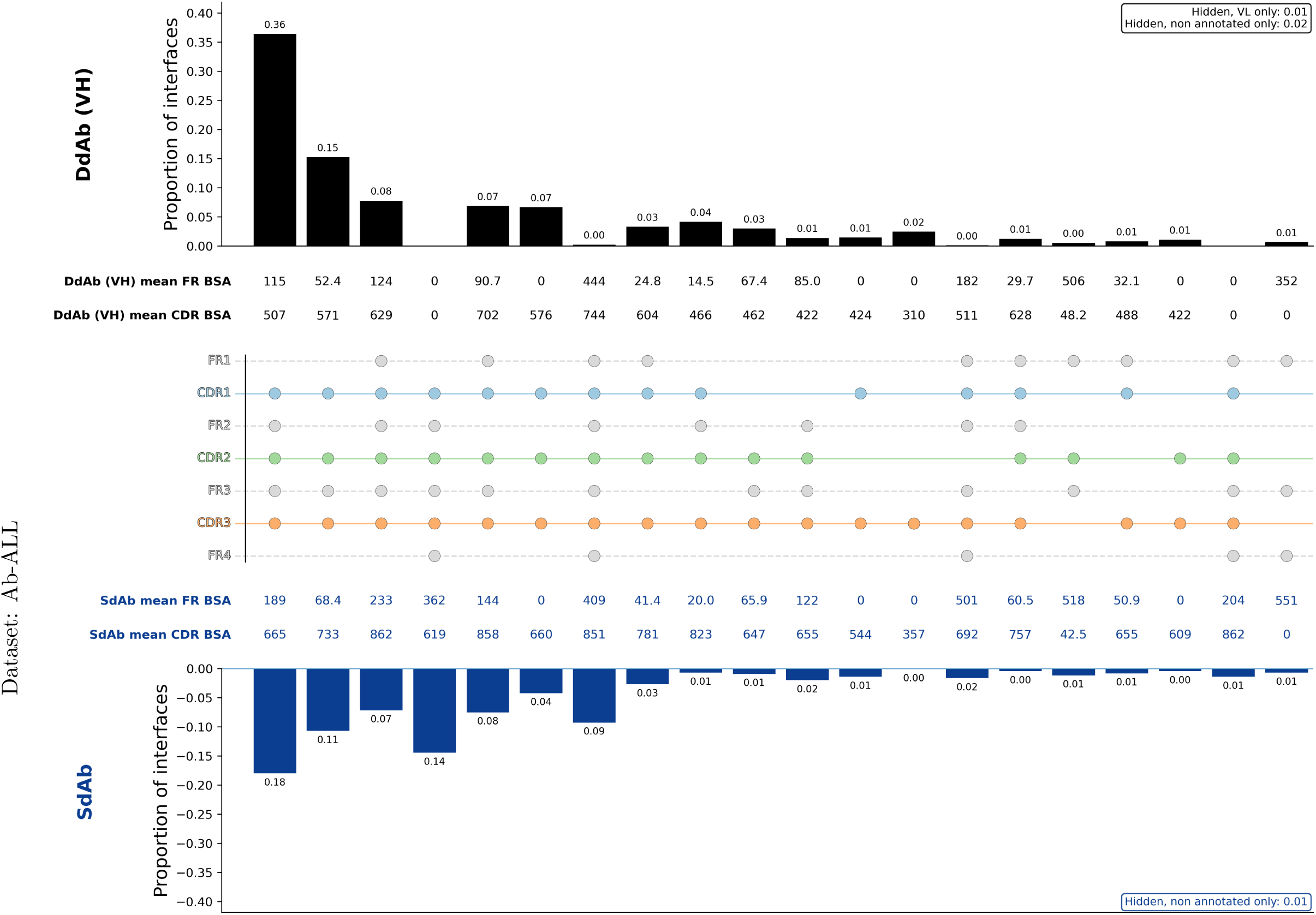
Contact profiles DdAb VH vs SdAb for the dataset Ab-ALL with the solvated model. A profile characterizes which regions of the heavy chain, out of CDRs and FRs, participate to the interface. The top 20 profiles (SdAb + DdAb) are displayed.

#### • SdAb and the CDR1+CDR2+CDR3 profile

For SdAb, previous work reported that ∼ 50% of cases display a pure CDR1+CDR2+CDR3 profile [14]; we obtain instead a fraction of 7 %. To understand this striking difference, we recompute profiles by imposing a minimum BSA of 40 Å^2^ per region to be considered in the footprint of the interface (Fig. S13). Although the agreement is not exact, partly due to a different dataset and differences in interface calculation methods, the profiles are in much better agreement. The thresholding from [14] is not ideal though. On the one hand, in Ab-ALL, SdAb have on average 934 Å^2^ of BSA so that the threshold used is ∼ 4%. On the other hand, using an energy density of ∼ 10 cal/mol per Å^2^ of buried surface area [41, 42], one can estimate the free energy error induced by the threshold to ∼ 0.4 *kcal/mol*, which is significant. (Recall that 1.4*kcal/mol* is a factor of 10 on the *K_d_*.)

#### • SdAb and prevalent profiles

Previous work reported that the two dominant profiles for SdAb are CDR1+CDR2+CDR3 (∼ 17%) and FR1+CDR1+FR2+CDR2+FR3+CDR3 (∼ 17%). (NB: FR4 is not used to define binding modes in [14].) For the former profile we get 4%, and for the latter 9 + 7% (with and without the FR4). With a 40Å^2^ threshold to consider a region part of the footprint, we found that, for SdAb, 3 footprints dominate the dataset: CDR1+FR2+CDR2+FR3+CDR3, CDR1+CDR2+CDR3 and CDR1+CDR2+FR3+CDR3 (with respectively 15%, 14 % and 11 % of the interfaces). For SdAb, using a threshold of 40Å^2^, despite some differences in footprint distributions compared with the Cambridge study, we recover the same overall trend: footprint types are more diverse than for DdAb. Although CDRs remain strongly represented at the interfaces, we also observe interfaces in which not all CDRs are involved, and where framework regions make a substantial contribution.

### All-residue profiles reveal greater footprint diversity in SdAb

In the sequel, we focus on the top 20 profiles (SdAb + DdAb VH) yielded by our model, so as to focus on recurrent footprints rather than isolated cases. Within this set, we retain for discussion profiles representing at least 5% of interfaces in either antibody class, and showing either a difference of at least 5 percentage points between SdAb and DdAb, near absence from one class, or a marked difference in framework-region contribution as assessed by FR BSA. We then distinguish profiles common to SdAb and DdAb from those enriched in either class. The corresponding analysis for the DdAb light chain is provided in Fig. S14.

- Common profiles:

**–** All CDRs + FR1-2-3 (8 % for DdAb and 7 % for SdAb)
**–** All CDRs + FR1+3 (7 % for DdAb and 8 % for SdAb)
**–** Other footprints with less than 5 % on both
- Profiles used mostly by DdAb:

– All CDRs + FR2+3 (36 % for DdAb vs 18 % for SdAb), with 115Å2 of mean FR BSA for DdAb and 189Å2 for SdAb.
– All CDRs + FR3 (15 % for DdAb vs 11 % for SdAb), with respectively 52Å2 vs 68Å2 mean FR BSA.
- Profiles mostly used by SdAb:

**–** All CDRs + FR2+3+4 ( 0 % for DdAb vs 14 % for SdAb) with more than half the BSA on FR regions for SdAb.
**–** All regions contact ( 0 % for DdAb vs 9 % for SdAb) with half the BSA on FR regions for SdAb.

Overall, SdAb display a broader diversity of interface footprints. In SdAb, three distinct interface footprint types account for 43% of the interfaces, whereas in DdAb, only one footprint type already account for 36% of the interfaces. For footprint types shared by SdAb and DdAb, framework regions also contribute more to the BSA in SdAb interfaces (Fig. 9).

**Figure 9:**
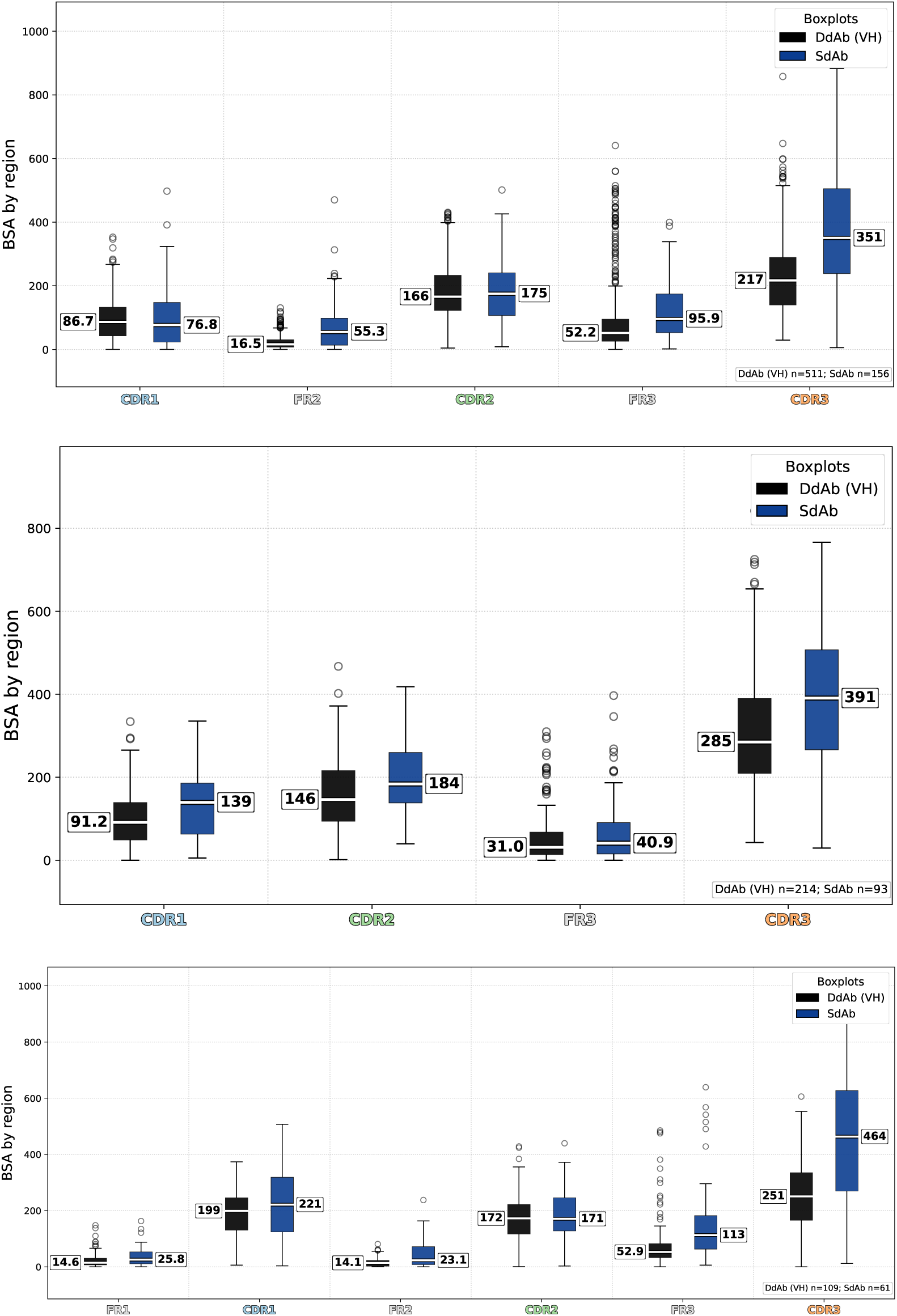
Dataset Ab-ALL, solvated model: Buried surface area (BSA) contribution per region for selected contact profiles. For identical interface footprints (Fig. 8), SdAbs show a larger framework contribution to BSA than DdAb VH interfaces. The three footprints were selected as the most prevalent across DdAb VH and SdAb interfaces, with non-zero representation in both groups.

These analysis point out the importance of framework regions in SdAb binding. Framework regions should thus be considered as full contributors to the binding interface, rather than as merely structural supports.

## 4 Discussion and outlook

### Geometry and binding modes

Using an integrated interface model, we analyze the commonalities and differences between interfaces made by SdAb and DdAb. In contrast to previous reports, we show that SdAb form interfaces with buried surface area comparable to that of DdAb, despite involving fewer residues and atoms. This ability stems from their more convex paratopes, whose atoms are more exposed to the binding partner than their counterparts in DdAb, thereby allowing each residue to establish more contacts. The longer CDR3 loops characteristic of SdAb follow this principle. Because these interfaces are tightly packed, the sum of the contact densities on the antibody and antigen sides remains essentially constant. We also note that while interfaces with SdAb tend to be more convex–which corresponds to the classical image of CDR inserted into a cleft or pocket, SdAb (resp. DdAb) also make concave (resp. convex) interfaces. There is no clear strict rule. Our analysis also stresses the role of FR and CDRs. We indeed show that FR regions play a role significantly more important than reported, both in terms of binding profiles and interface packing. FR regions not only have an architectural role, but significantly contribute to binding. This is expected as a consequence of the absence of a VL domain imposing steric hindrance, thereby allowing SdAb FRs to fulfill roles beyond dimerization.

Let us now bridge the gap between these structural observations and the thermodynamics and kinetics of binding.

### Thermodynamics implications

As recalled in Introduction, the thermodynamics of binding are complex, a fact which is acknowledged by affinity prediction models. In particular, regressing *K_d_* using the BSA yields a global trend stating the affinity increases when BSA increases [42], yet the correlation remains poor. Predictions improve with linear models taking into account conformational changes upon binding [43], or even better atomic packing properties [32], yet, such models require high-resolution bound and unbound crystal structures.

Nevertheless, the insights on the geometry of interfaces yielded by our models remain of interest to perform a qualitative analysis. We have seen that both types of interfaces have similar BSA and a comparable total contact density. In other words, the convex interfaces of SdAb concentrate the same contact network over fewer SdAb residues, while distributing them over more antigen residues. This uneven distribution of contacts calls for the following comments. First, the binding energy may be more concentrated on the SdAb side, with highly connected a.a. acting as hot spots *i.e.* residues yielding a significant ΔΔ*G* change (say above 2 kcal/mol), whereas energetic contributions may be more distributed over the antigen side. Second, a higher coordination for SdAb residue can improve the enthalpic efficiency per residue, via wan der Waals contact, hydrogen bonds, or electrostatic interactions. This enhancement can counterbalance a higher entropic penalty upon binding, since, unless pre-conditioning is at play [4, 5], long CDR3 may lose substantial conformational entropy. The presence of hot spots characterized by these geometric criteria is also consistent with classical analysis stating that for small interfaces (say BSA *<* 2000Å^2^), the energy density decreases linearly [42]. In other words, the burial of 1Å^2^ of surface area is more energetically favorable for smaller interfaces, with values dropping from ∼ 13 cal/mol/Å^2^ (at BSA = 500Å^2^) down to ∼ 4cal/mol/Å^2^ (at BSA = 200Å^2^) [42]. Hot spots with large ΔΔ*G* are sensitive to mutagenesis, both in nature and for engineering. These a.a. are of high interest to tune the binding affinity and specificity by performing a small number of pointwise mutations.

Finally, the desolvation patterns are expected to differ. Exposed atoms from SdAb in convex regions are highly solvent exposed before association, and their burial can be favorable if they are hydrophobic or if binding releases constrained waters gaining entropy in the bulk. Conversely, burying exposed polar groups is costly unless their hydrogen bonding potential is satisfied. A complementary concave antigen region can also contain structural water. Expelling it can be favorable, but retaining cavities or unsatisfied polar atoms can be unfavorable.

Summarizing, one expects flat interfaces to display a moderate enthalpy gain and a small entropy penalty, and convex interfaces to harbor a larger enthalpy gain and a larger entropy penalty, possibly focused on few residues. Moreover, depending on the nature of exposed amino acids, curvature can change the enthalpy-entropy balance via solvation effects, without substantially changing the total BSA. Both observations are of interest in mutagenesis studies, as they may help identify residues to be mutated in priority. Indeed, our analysis hints at a selection of candidates a.a. based on geometric criteria involving the curvature the molecular surface and its solvent accessibility.

### Towards quantitative predictions

The previous qualitative analysis naturally raises the question of quantitative predictions. From the structural standpoint, epitope and binding modes prediction remain challenging in the era of AI [44, 45]. The problems are even more complex when it comes to thermodynamics and kinetics. Although the theory of free energy in non-covalent binding is well established [46], the relevant molecular motions occur on time scales of milliseconds and beyond [47], making routine molecular dynamics simulations impractical at this stage. The thermodynamic complexity is also a clear difficulty for current AI based co-folding algorithms overlooking entropic effects and the integration of water molecules at the interface. Kinetics can also play a key role, as delivery in specific organs *e.g.* the brain may depend on the dissociation kinetics [48].

Two specific research veins are worth investigating to overcome these difficulties. On the one hand, in the wake of AlphaFold2 [49], deep learning approaches may eventually enable the learning of thermodynamics and kinetics directly from structural data. But current prediction frameworks based on flow matching and diffusion models face various challenges, including the need of large molecular dynamics datasets for training, the question of sampling coherently the main chain and the side chains, and the control of potential biases incurred during the learning phase. In parallel to these works, geometry and physics informed methods offer an appealing strategy to handle efficiently the case of antibodies and their CDRs. Following the seminal work on kinematic loop closure [50], advanced kinematic models for backbones [51, 52] combined with statistical models on side-chains [53] now enable the simulation of long flexible loops. These methods offer a promising route to efficiently explore the conformational landscapes of CDRs in particular and predict their thermodynamic properties. We note that such explorations should take into account the secondary structure contents of CDRs [44], a topic for which generative models have been recently developed [54].

Progress along either of these directions would undoubtedly accelerate the design of nanobodies and further unlock their therapeutic potential.

## 5 Artwork

**Table 1:**
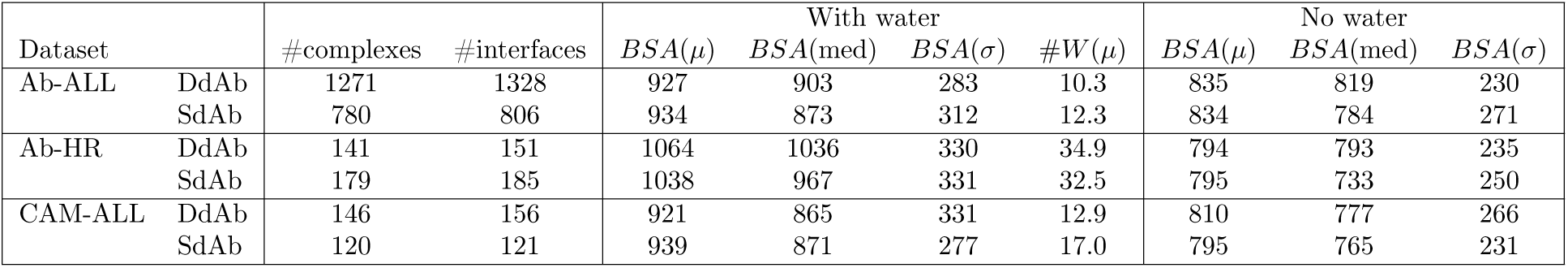
Interfaces, global statistics, with and without crystallographic water molecules. Ab-ALL: all structures with resolution 4 or better; Ab-HR: high resolution structures; CAM-ALL: Cambridge dataset [14, 13]. NC: number of complexes; NI: number of interfaces. BSA: buried surface area; #W: mean number of water molecules included in the interface. W/nw: with and without water molecules included in the interface calculation.

