## Supporting Information for "Distinct geometries, comparable interfaces: binding modes and thermodynamic implications in conventional and single domain antibodies"

### S1 Supporting Information

#### S1.1 Dataset curation, composition and target diversity

##### S1.1.1 De-duplication strategy

Previous studies used sequence-based de-duplication strategies. Mitchell *et al.* removed perfect SdAb duplicates to define the CAM-ALL dataset [13, 14], whereas Gordon *et al.* used a 95% identity cutoff across CDR residues, retaining selected complexes when they bound substantially different antigens [15]. Although CDR sequence-based de-duplication efficiently limits sequence redundancy, our analyses focus primarily on antibody–antigen interfaces, which may differ even for identical partners studied under different pH or ionic-strength conditions. We therefore refined the filtering procedure to preserve sequence-similar antibodies whenever they form distinct interfaces. We used a hierarchical de-duplication procedure in which targets were first grouped by sequence similarity and antibodies within each target cluster were then grouped by sequence similarity across their IMGT-defined CDR residues. Within each antibody cluster, antibody-side Voronoi interfaces were encoded as *interface strings*, using the one-letter amino-acid code for interfacial residues and dashes for non-interfacial positions. After alignment, normalized Hamming distances were computed between interface strings. Complexes differing by less than 5% were considered redundant, and only the highest-resolution structure was retained from each sub-cluster.

The final list of complexes used is provided in the file http://sbl.inria.fr/data/SdAb-DdAb/SdAb-DdAb-datasets.csv.

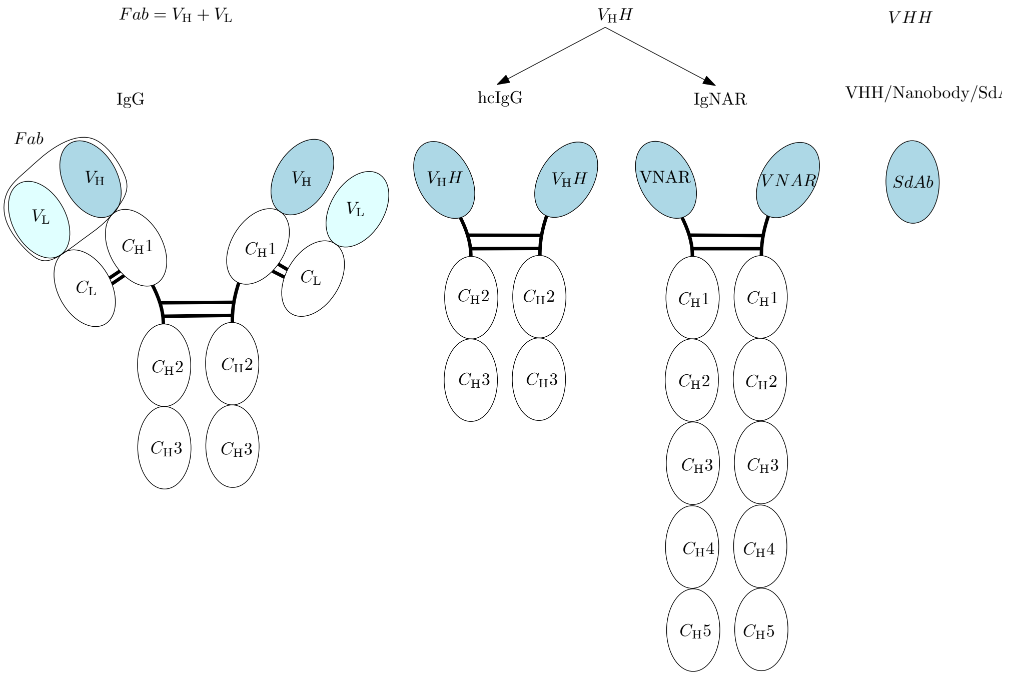


Figure S1: **Variable domains studied in this work–blue regions.** We consider VDs found in IgG and IgD, namely $Fab = V_H + V_L$, $V_HH$ found in heavy-chain-only antibodies (VHH in camelids and VNAR in cartilaginous fish), and single-domain antibodies

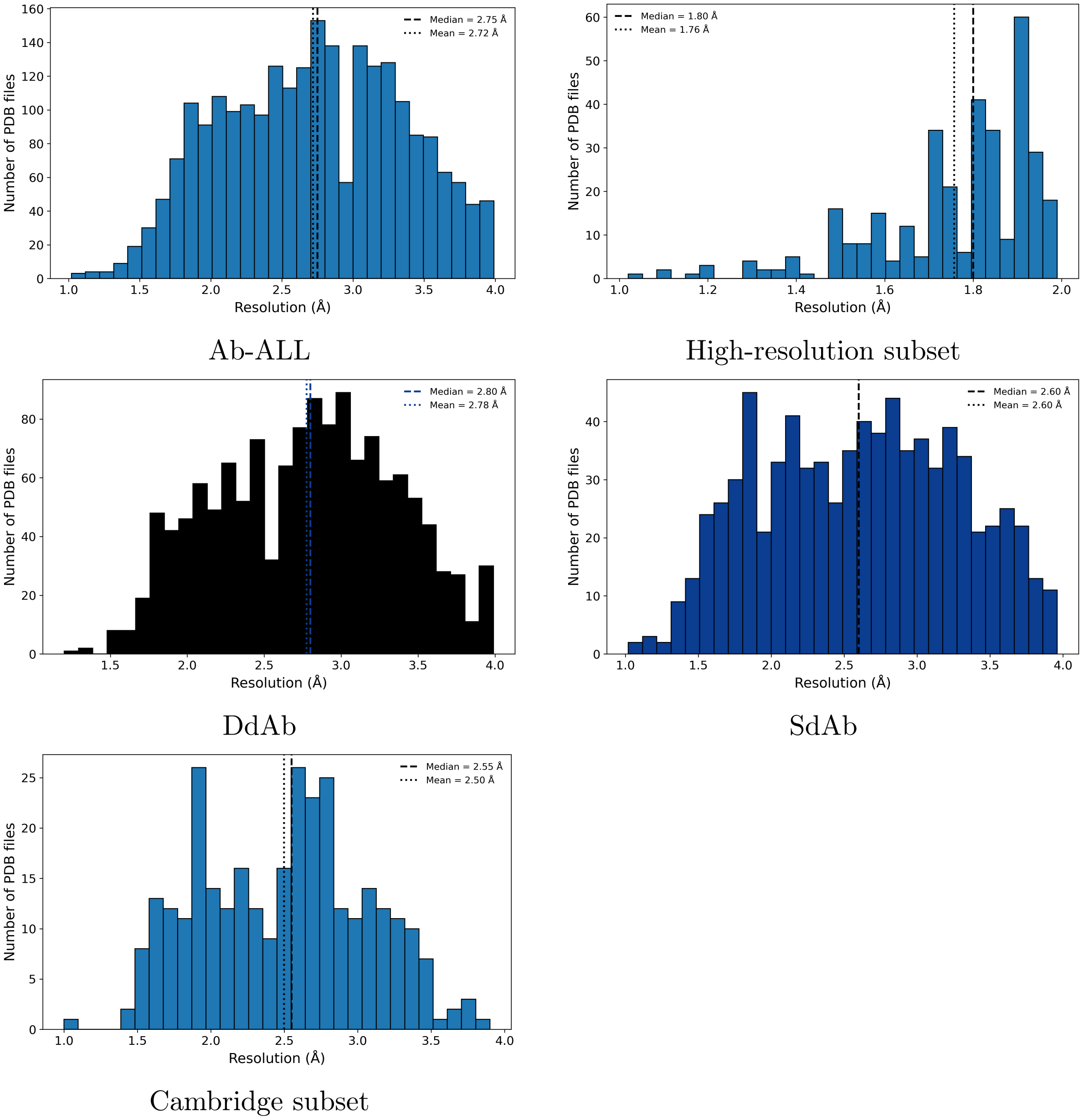


Figure S2: **Resolution distributions for the structural datasets.** Histograms of PDB resolution for DdAb and SdAb separately, and for the high-resolution and Cambridge subsets used in the analysis.

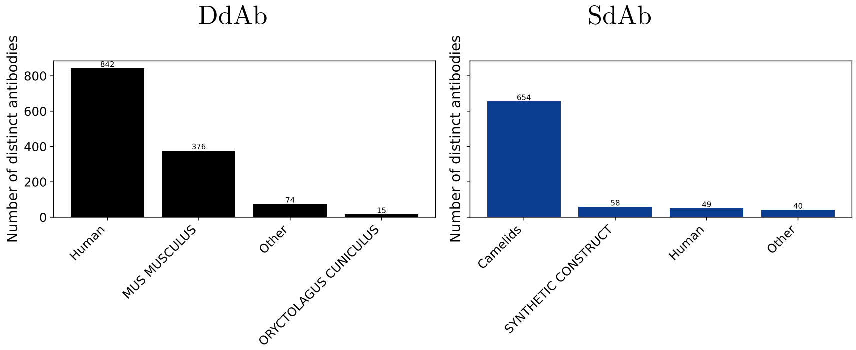


Figure S3: **Species distribution of DdAb and SdAb in the dataset.**

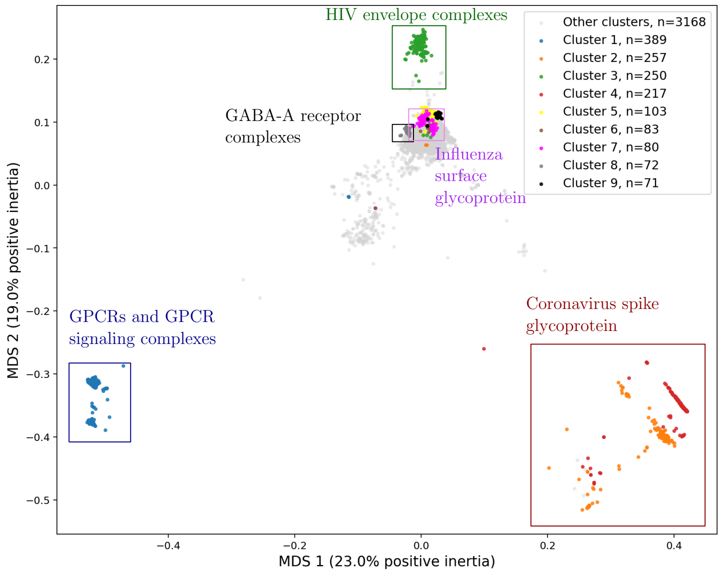


**(A) All antibody complexes**

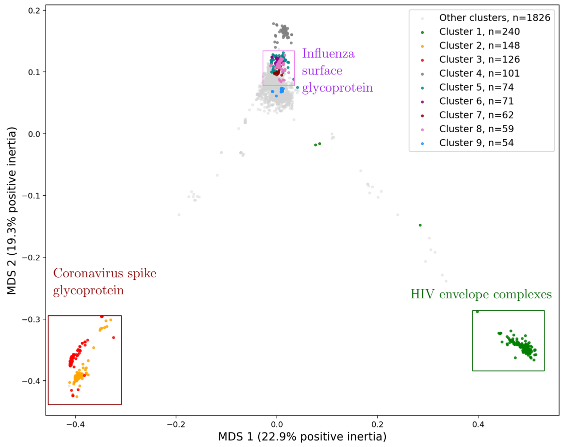


**(B) DdAb**

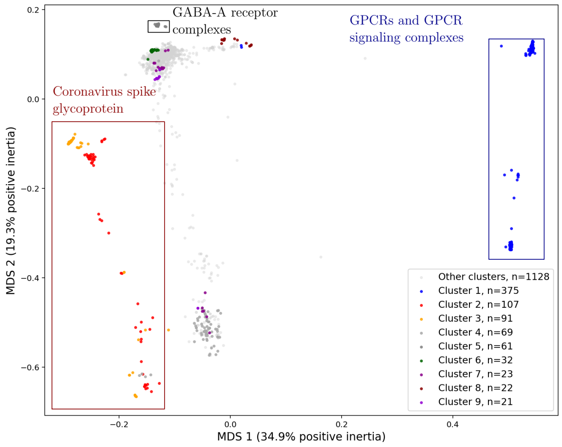


**(C) SdAb**

Figure S4: **2D embedding of targets using sequence distances and multi-dimensional scaling (MDS).** All sequences of targets in databases are processed (IMGT/3Dstructure-DBand SAbDab-nano) – one point per target. Colors indicate the largest clusters found by cutting the dendrogram at a fixed distance threshold (see main text); smaller clusters are grouped and displayed in gray. **(A)** All antibody formats *i.e.* SdAb and DdAb. **(B)** DdAb. **(C)** SdAb. Shared target families are observed across antibody formats, including spike proteins, while other clusters appear more format-specific, such as HIV envelope glycoproteins in DdAb and adrenergic receptors in SdAb.

| Target family | AB | DdAb | SdAb | Interpretation |
|---|---|---|---|---|
| Coronavirus spike glycoprotein | Clusters 2 and 4, large clusters | Clusters 2 and 3 | Clusters 2 and 3 | Shared viral target family. The presence in all three analyses likely reflects the strong structural sampling of coronavirus spike, with possible splitting according to construct, chain composition, or oligomeric state. |
| HIV envelope gp120/CD4 complexes | Cluster 3 | Cluster 1 | Not among the main displayed clusters | Coherent DdAb-enriched viral glycoprotein cluster. This likely reflects conventional antibody structures against HIV envelope complexes rather than a general target class shared across formats. |
| Influenza surface glycoproteins | Cluster 7 | Clusters 5 and 8 | Influenza polymerase-related cluster, not the same surface glycoprotein family | Mostly DdAb in the displayed clusters. This suggests that conventional antibodies dominate the influenza HA/NA target landscape, whereas SdAb examples may correspond to different influenza proteins. |
| Lysozyme | Cluster 6 | Cluster 7 | Cluster 9 | Shared historical model antigen. Its recurrence in both formats indicates a dataset composition effect rather than a format-specific biological preference. |
| GPCRs and GPCR signalling complexes | Cluster 1 | Not among the main displayed clusters | Clusters 1 and 8 | Mostly SdAb-associated in the displayed clusters. This is biologically plausible because nanobodies are often used to stabilize GPCR active states or GPCR signalling complexes. Some entries include fusion partners, so this cluster should be interpreted as construct-dependent. |
| GABA-A receptor complexes | Cluster 8 | Not clearly isolated among the top displayed DdAb clusters | Cluster 5 | Membrane receptor family represented in both global and SdAb views. |
| KcsA potassium channel | Cluster 9 | Cluster 6 | Not among the main displayed clusters | Coherent DdAb-associated membrane protein cluster in the displayed representatives. This likely reflects repeated structures against the same bacterial channel target. |
| Integrin $\alpha$IIb/$\beta$3 | Not among the main displayed AB clusters | Cluster 9 | Not among the main displayed clusters | DdAb-specific cluster in the displayed representatives |
| Ricin and GFP | Not among the main displayed AB clusters | Not among the main displayed clusters | Clusters 6 and 7 | Small but biologically coherent SdAb-enriched clusters. They likely reflect recurrent nanobody benchmark or tool-antigen systems. |

Table S1: Main biologically interpretable target-sequence clusters observed after hierarchical clustering at distance threshold 0.6. The corresponding MDS visualizations are shown in Fig. S4, with the format-specific analyses shown for DdAb in panel B and SdAb in panel C. Cluster numbers are specific to each analysis because clusters were renumbered by size within each dataset.

#### S1.2 Interface model

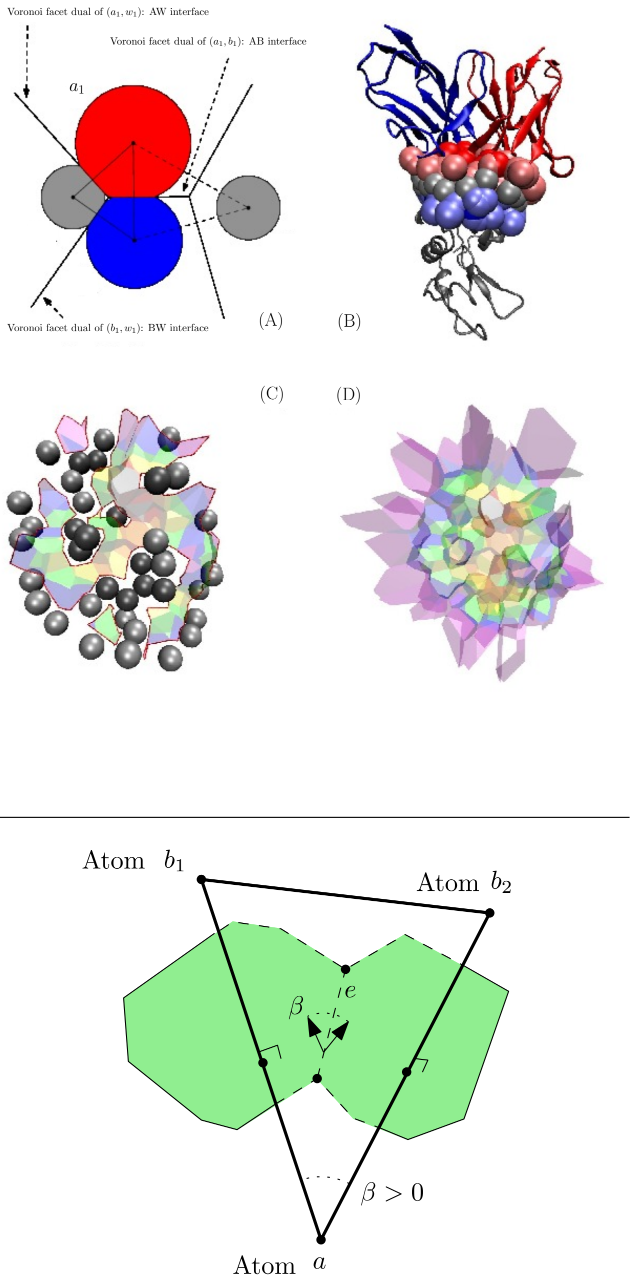


Figure S5: **The Voronoi interface model and its curvature. (Top) (A)** 2D model **(B) Antibody:antigen** Model: 1vfb.pdb **(C)** AB interface and interfacial water **(D)** ABW interface. **(Bottom)** Atoms represented as points rather than balls to avoid cluttering. The atom $a$ from partner $A$ is in contact with two atoms $b_1, b_2$ from partner $B$. The edges $ab_1$ and $ab_2$, from the weighted Delaunay triangulation/$\alpha$-complex define the contacts. The dual Voronoi facets in green define the $AB$ interface which separates partners. The local curvature of the interface is length$(e) \times \beta$. (NB: this quantity is akin to a mean curvature and not a gaussian curvature.) The curvature is positive (resp. negative) when the interface observed from partner $A$ is convex (resp. concave). The AW-BW interface is defined similarly: it involves a water molecule $w$ sandwiched between two atoms $a$ and $b$ from the partners $A$ and $B$.

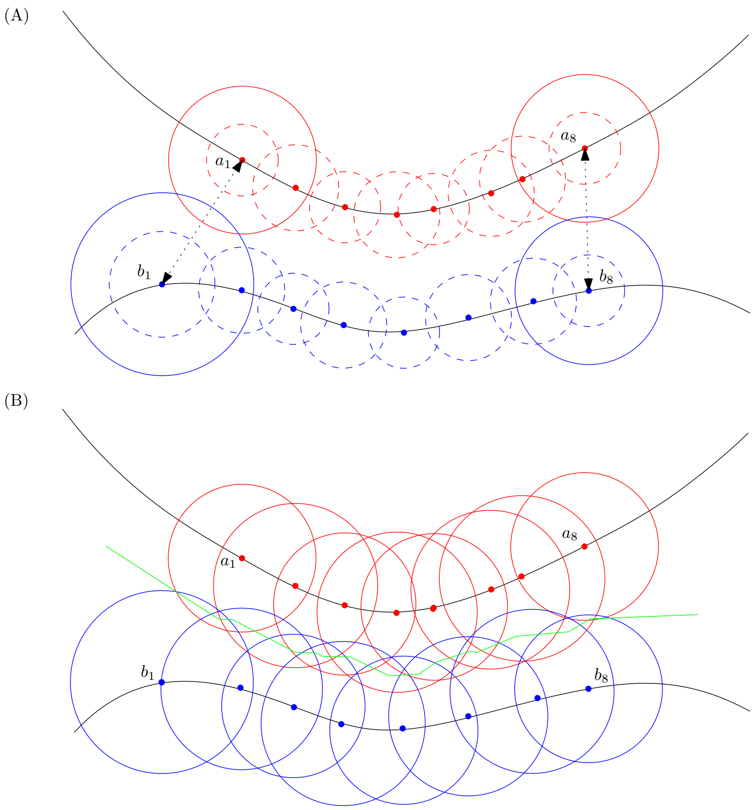


Figure S6: **Interface models: distance based versus Voronoi.** Atomic representations: dashed (resp. solid) for the van der Waals (resp. solvent accessible) representation. **(A)** Distance-based. Two atoms whose SAS intersect may be at a distance larger than the threshold used to define the contacts (typically $r = 4.5\text{Å}$), notably on the interface rim, *e.g.* for the pairs $(a_1, b_1)$ and $(a_8, b_8)$. **(B)** Voronoi interface. The Voronoi interface instead captures all atoms with buried surface area.

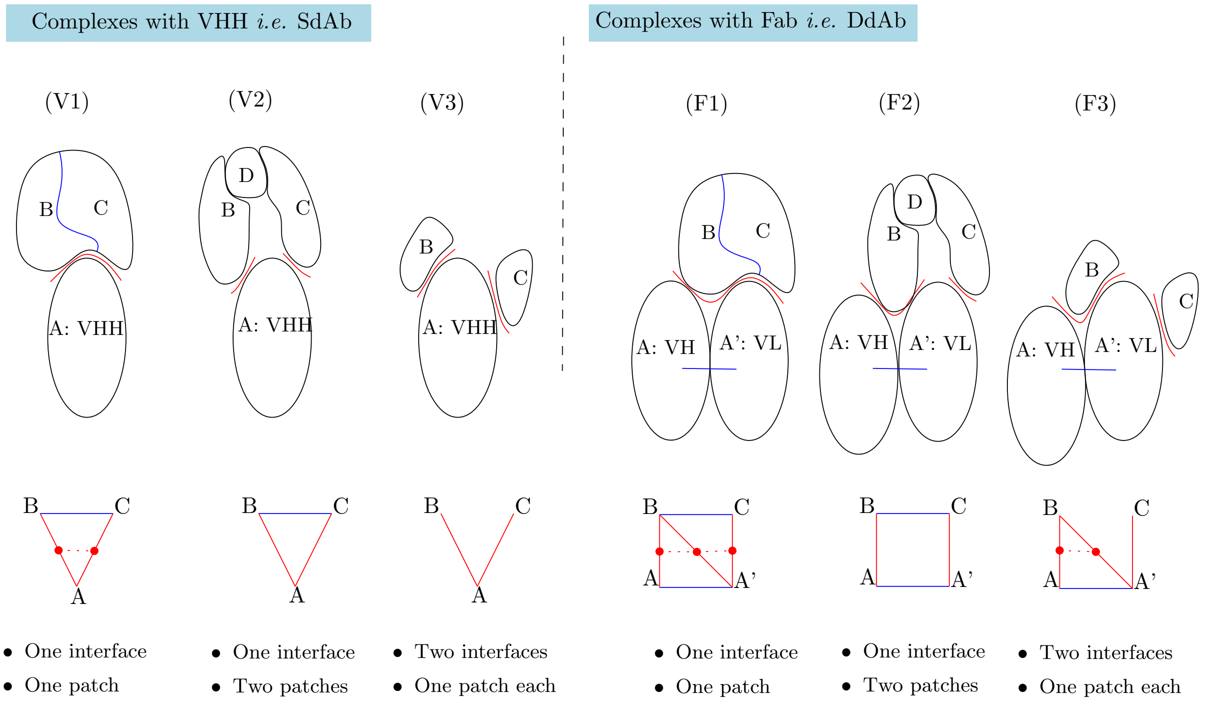


Figure S7: **Automating the study of interfaces between antibodies and antigens.** A SdAb (resp. DdAb) consists of a single (resp. two) variable domain(s). The antigen found in the asymmetric unit of a crystal structure may consist of several chains. The number of **connected components** of these chains defines the number of interfaces. **(A)** Geometry of the subunits. **(B)** Graph depicting contacts between subunits. The number of connected components of each row defines the number of partners, whence the number of interfaces. The red vertical/slanted segments define binding patches, which may be connected or not (red horizontal dashed segments).

#### S1.3 Parsing and annotation

**Antibody parsing, annotation and interface computation.** All structures were processed through a harmonized antibody-complex loading pipeline. For each entry, SEQRES sequences and experimentally observed structural sequences were extracted when available. Variable domains were annotated in the IMGT scheme [24], using database-provided annotations whenever possible and `anarcii` numbering otherwise [56]. The annotation was assigned at the sequence level and transferred to the observed structural sequence by alignment when required.

Antibody chains were classified as heavy or light chains using chain names, metadata, IMGT descriptions, sequence motifs and, when needed, IMGT numbering. Antibody type was inferred from species, naming conventions and chain composition. For comparative analyses, paired heavy–light variable-domain antibodies were grouped as DdAb, whereas heavy-chain-only antibodies, nanobody-like entries, VHHs and VNARs were grouped as SdAb.

**Antibody types and their inference.** The antibody type is inferred using a rule-based procedure combining species information, antibody naming conventions, and chain composition.

First, the species of origin is examined. Antibodies originating from specific shark species (*Squalus acanthias*, *Orectolobus maculatus*, or *Ginglymostoma cirratum*) are classified as `VNAR` when they contain heavy chains but no light chains. If the species corresponds to a camelid (species name containing markers such as *Lama*, *Camelidae*, *Camelus*, *Vicugna*, or *Llama*) and the antibody contains heavy chains but no light chains, it is classified as `HC_IGG`. If species-based rules do not apply, the antibody name and title are inspected. If they contain keywords such as *nanobody*, *vhh*, *sdab*, or *vnar*, the antibody is classified as `NANOBODY`.

Finally, if generic antibody-related terms are detected (e.g., *antibody*, *mAb*, *Fab*, *IgG*, or *immunoglobulin*), the chain composition is used: antibodies containing heavy chains but no light chains are classified as `HC_IGG`, while antibodies containing both heavy and light chains are classified as `DDAB`. If none of these rules apply, the antibody is labeled as `GENERIC` (or `NANOBODY` if the pdb is from SAbDab)

| Object | Group | Values | Meaning |
|---|---|---|---|
| AntibodyType | DdAb | DDAB | Classical double domain antibody interface, involving paired heavy and light variable domains. |
| | SdAb | VHH, HC_IGG, VNAR, NANOBODY | Single domain antibody classes, grouped together for comparisons against DdAbs. |
| | Generic | GENERIC | Generic type, used when no specific class is assigned. Not used in the analysis |
| Ig_region_type | Framework regions | FR1, FR2, FR3, FR4 | Conserved framework regions of the variable domain. |
| | CDR regions | CDR1, CDR2, CDR3 | Complementarity determining regions, usually enriched in target binding residues. |
| | Unannotated | SEQ | Residues present in the sequence or structure but not assigned to an IMGT FR or CDR region. |
| AtomicContactType | Direct contacts | AB | Direct antibody, target atomic contact. |
| | Water contacts | AW | Direct antibody, water atomic contact. |
| | Water contacts | BW | Direct target, water atomic contact. |
| WaterBridgedContact | Water mediated contact | AW - BW | A water bridged contact is defined when an antibody atom and a target atom contact the same water molecule. |

Table S2: **Main categorical types used to describe antibody, region and atomic contact annotations.** `AntibodyType` values are grouped into generic, DdAb and SdAb categories for downstream comparisons. `Ig_region_type` follows the IMGT subdivision into framework and CDR regions, while `SEQ` is kept separate because it corresponds to residues that could not be assigned to a specific annotated region. Atomic contacts are represented either as direct contacts, `AB`, `AW` or `BW`, or as water mediated triplets when `AW` and `BW` involve the same water molecule.

#### S1.4 Variable-domain and interface statistics

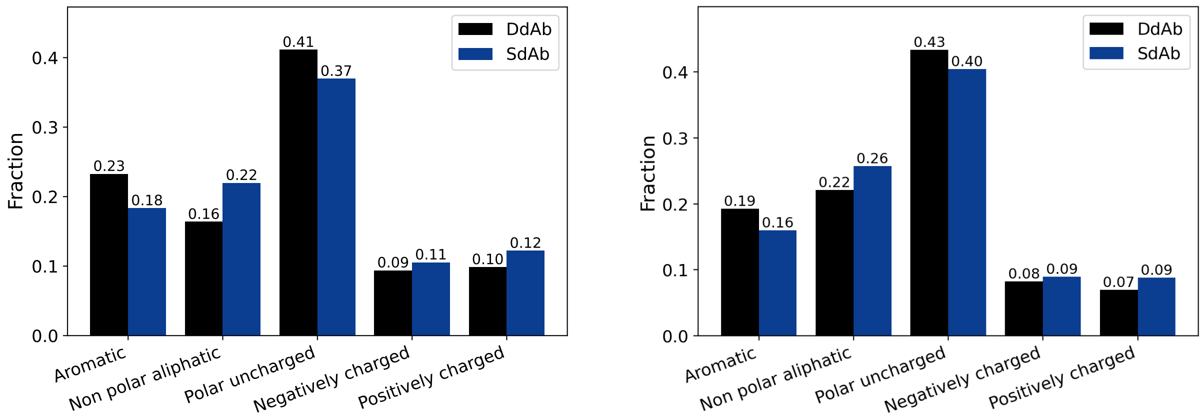


**(A)** Interface residues

**(B)** CDR residues

Figure S8: **Dataset Ab-ALL, solvated model: Amino-acid category distributions in SdAb and DdAb interfaces and CDRs.** (A) Amino-acid category distribution among interface residues. (B) Amino-acid category distribution among CDR residues.

Table S3: **Amino-acid enrichment by antibody group and region using mean frequencies. Dataset: all VD.** For each amino acid, enrichment is $[p/(1-p)]/[p_{\text{ref}}/(1-p_{\text{ref}})]$, where $p_{\text{ref}}$ is the corresponding frequency in the same region of the combined DDAB+SDAB dataset. Values above 1 indicate enrichment and values below 1 indicate depletion. Green cells are enriched, red cells are depleted, and color intensity is proportional to $|\log_2(E)|$. $N$ is the number of usable domains in the displayed group. These counts can vary between regions because domains with a missing, length-inconsistent or empty regional annotation are excluded.

| | | | **Aromatic** | | | **Non polar aliphatic** | | | | | | **Polar uncharged** | | | | | | **Charged (-)** | | **Charged (+)** | | |
|---|---|---|---|---|---|---|---|---|---|---|---|---|---|---|---|---|---|---|---|---|---|---|
| **Group** | **Region** | $N$ | **F** | **W** | **Y** | **P** | **M** | **I** | **L** | **V** | **A** | **G** | **S** | **T** | **N** | **Q** | **C** | **D** | **E** | **H** | **K** | **R** |
| DDAB | Total | 2909 | 1.04 | 1.01 | 1.00 | 1.06 | 0.94 | 1.10 | 1.02 | 0.95 | 0.93 | 0.97 | 1.05 | 1.04 | 0.91 | 1.00 | 0.97 | 0.99 | 0.96 | 1.07 | 1.03 | 0.92 |
| DDAB | CDRs | 2908 | 1.01 | 1.03 | 1.05 | 1.03 | 0.98 | 0.93 | 0.95 | 0.98 | 0.92 | 0.94 | 1.05 | 0.97 | 1.05 | 1.24 | 0.76 | 1.00 | 0.87 | 1.03 | 0.96 | 0.87 |
| DDAB | FRs | 2909 | 1.04 | 1.01 | 0.98 | 1.07 | 0.93 | 1.17 | 1.03 | 0.94 | 0.93 | 0.97 | 1.06 | 1.07 | 0.83 | 0.96 | 0.98 | 0.99 | 0.97 | 1.13 | 1.03 | 0.92 |
| DDAB | FR1 | 2909 | 0.91 | 1.01 | 1.22 | 1.16 | 1.26 | 1.28 | 0.86 | 1.01 | 0.94 | 0.78 | 1.04 | 1.25 | 1.10 | 0.92 | 0.99 | 1.22 | 0.98 | 0.99 | 1.27 | 0.96 |
| DDAB | CDR1 | 2908 | 0.92 | 1.10 | 1.08 | 0.53 | 0.60 | 0.99 | 0.87 | 1.02 | 0.80 | 0.98 | 1.10 | 0.94 | 1.11 | 1.28 | 0.60 | 0.97 | 0.78 | 0.93 | 1.06 | 0.64 |
| DDAB | FR2 | 2908 | 0.38 | 1.08 | 1.17 | 1.09 | 0.87 | 1.24 | 1.24 | 0.92 | 0.77 | 0.96 | 1.04 | 0.97 | 1.12 | 1.11 | 0.31 | 0.78 | 0.70 | 1.22 | 1.08 | 0.74 |
| DDAB | CDR2 | 2908 | 1.06 | 0.84 | 1.19 | 1.13 | 0.78 | 0.86 | 0.91 | 1.03 | 1.24 | 0.88 | 1.03 | 0.84 | 1.09 | 0.98 | 0.47 | 1.07 | 1.07 | 0.83 | 1.16 | 0.71 |
| DDAB | FR3 | 2908 | 1.11 | 0.89 | 0.93 | 1.01 | 0.82 | 1.04 | 1.02 | 0.88 | 0.99 | 1.15 | 1.13 | 1.04 | 0.78 | 0.99 | 1.00 | 0.96 | 1.05 | 0.96 | 0.78 | 1.00 |
| DDAB | CDR3 | 2908 | 1.11 | 1.09 | 0.99 | 1.05 | 1.08 | 0.89 | 0.97 | 0.94 | 0.84 | 0.95 | 1.03 | 1.08 | 0.92 | 1.24 | 0.82 | 0.99 | 0.87 | 1.09 | 0.75 | 0.97 |
| DDAB | FR4 | 2907 | 1.31 | 0.86 | 0.78 | 0.99 | 1.21 | 1.27 | 1.26 | 0.89 | 0.97 | 1.06 | 0.81 | 0.97 | 0.85 | 0.71 | 0.28 | 1.23 | 1.26 | 0.78 | 1.24 | 0.92 |
| SDAB | Total | 882 | 0.88 | 0.96 | 0.99 | 0.81 | 1.19 | 0.69 | 0.93 | 1.18 | 1.23 | 1.10 | 0.83 | 0.87 | 1.31 | 0.98 | 1.09 | 1.02 | 1.12 | 0.78 | 0.91 | 1.28 |
| SDAB | CDRs | 878 | 0.96 | 0.89 | 0.84 | 0.91 | 1.07 | 1.25 | 1.16 | 1.07 | 1.26 | 1.19 | 0.83 | 1.11 | 0.84 | 0.25 | 1.79 | 1.01 | 1.43 | 0.90 | 1.14 | 1.43 |
| SDAB | FRs | 882 | 0.86 | 0.95 | 1.06 | 0.77 | 1.24 | 0.46 | 0.92 | 1.21 | 1.22 | 1.09 | 0.81 | 0.79 | 1.58 | 1.15 | 1.08 | 1.02 | 1.10 | 0.57 | 0.91 | 1.25 |
| SDAB | FR1 | 882 | 1.30 | 0.97 | 0.29 | 0.50 | 0.15 | 0.10 | 1.49 | 0.98 | 1.20 | 1.80 | 0.88 | 0.23 | 0.68 | 1.27 | 1.05 | 0.29 | 1.05 | 1.02 | 0.14 | 1.12 |
| SDAB | CDR1 | 878 | 1.26 | 0.68 | 0.75 | 2.60 | 2.33 | 1.04 | 1.44 | 0.95 | 1.69 | 1.06 | 0.69 | 1.19 | 0.64 | 0.12 | 2.33 | 1.08 | 1.73 | 1.23 | 0.81 | 2.23 |
| SDAB | FR2 | 878 | 3.15 | 0.74 | 0.46 | 0.70 | 1.43 | 0.25 | 0.27 | 1.26 | 1.81 | 1.14 | 0.88 | 1.10 | 0.59 | 0.65 | 3.30 | 1.74 | 2.06 | 0.27 | 0.74 | 1.91 |
| SDAB | CDR2 | 851 | 0.79 | 1.55 | 0.39 | 0.55 | 1.76 | 1.52 | 1.33 | 0.90 | 0.27 | 1.43 | 0.89 | 1.61 | 0.71 | 1.08 | 2.81 | 0.75 | 0.77 | 1.58 | 0.45 | 2.02 |
| SDAB | FR3 | 878 | 0.63 | 1.35 | 1.24 | 0.96 | 1.62 | 0.87 | 0.94 | 1.40 | 1.02 | 0.51 | 0.59 | 0.86 | 1.77 | 1.02 | 1.00 | 1.13 | 0.83 | 1.13 | 1.76 | 1.01 |
| SDAB | CDR3 | 878 | 0.63 | 0.71 | 1.04 | 0.84 | 0.73 | 1.36 | 1.11 | 1.20 | 1.57 | 1.18 | 0.90 | 0.76 | 1.26 | 0.26 | 1.59 | 1.04 | 1.44 | 0.71 | 1.84 | 1.09 |
| SDAB | FR4 | 878 | 0.01 | 1.47 | 1.72 | 1.04 | 0.31 | 0.15 | 0.20 | 1.38 | 1.10 | 0.81 | 1.69 | 1.09 | 1.48 | 2.08 | 3.40 | 0.23 | 0.17 | 1.72 | 0.24 | 1.26 |

Table S4: **Amino-acid enrichment at the antibody-target interface using mean frequencies. Dataset: Ab-ALL; solvated model.** For each amino acid, enrichment is $[p/(1-p)]/[p_{\text{ref}}/(1-p_{\text{ref}})]$, where $p_{\text{ref}}$ is the corresponding frequency in the combined DDAB+SDAB dataset. Values above 1 indicate enrichment and values below 1 indicate depletion. Green cells are enriched, red cells are depleted, and color intensity is proportional to $|\log_2(E)|$. $N$ is the number of usable interfaces in the displayed group.

| | | | Aromatic | | | Non polar aliphatic | | | | | | Polar uncharged | | | | | | Charged (-) | | Charged (+) | | |
|---|---|---|---|---|---|---|---|---|---|---|---|---|---|---|---|---|---|---|---|---|---|---|
| **Group** | **Region** | $N$ | **F** | **W** | **Y** | **P** | **M** | **I** | **L** | **V** | **A** | **G** | **S** | **T** | **N** | **Q** | **C** | **D** | **E** | **H** | **K** | **R** |
| DDAB | Interface | 1393 | 0.97 | 1.02 | 1.16 | 1.00 | 0.78 | 0.96 | 0.90 | 0.86 | 0.77 | 1.01 | 1.14 | 0.98 | 1.09 | 0.84 | 0.69 | 1.01 | 0.84 | 1.15 | 0.98 | 0.82 |
| SDAB | Interface | 850 | 1.05 | 0.97 | 0.75 | 1.00 | 1.37 | 1.06 | 1.16 | 1.23 | 1.38 | 0.98 | 0.78 | 1.04 | 0.85 | 1.27 | 1.51 | 0.98 | 1.26 | 0.75 | 1.04 | 1.31 |

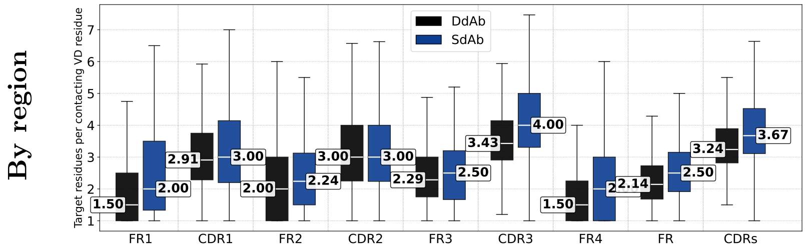


Figure S9: **Local contact density–using the ABW model which includes water-mediated contacts.** Refer to Fig. S5 for the definition of contacts. Local contact density is computed separately for each region. For each region, we first select the VD residues assigned to this region that actually contact the target, either directly or through a water molecule. We then compute the average number of target residues contacted by each of these contacting VD residues. The local contact density is higher for SdAb interfaces overall.

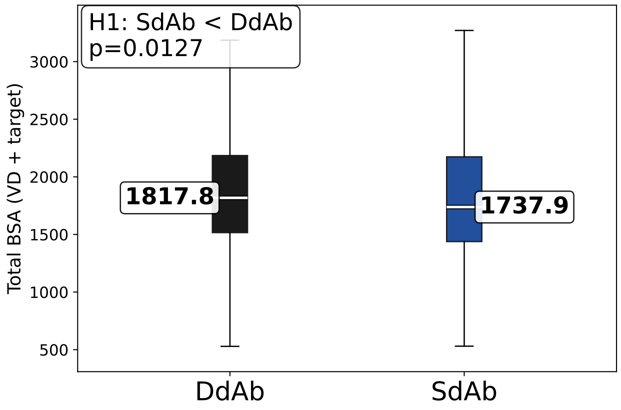


Figure S10: **Summed buried surface area on the variable-domain and target sides of the interface.** The boxplot shows the distribution of the total buried surface area obtained by summing the BSA measured on the antibody variable-domain side and on the target side of each interface, using the Ab-ALL dataset and the solvated interface model. This representation complements the VD-side BSA analysis by showing that, when both sides of the interface are considered together, SdAb and DdAb exhibit even more similar total buried surface areas.

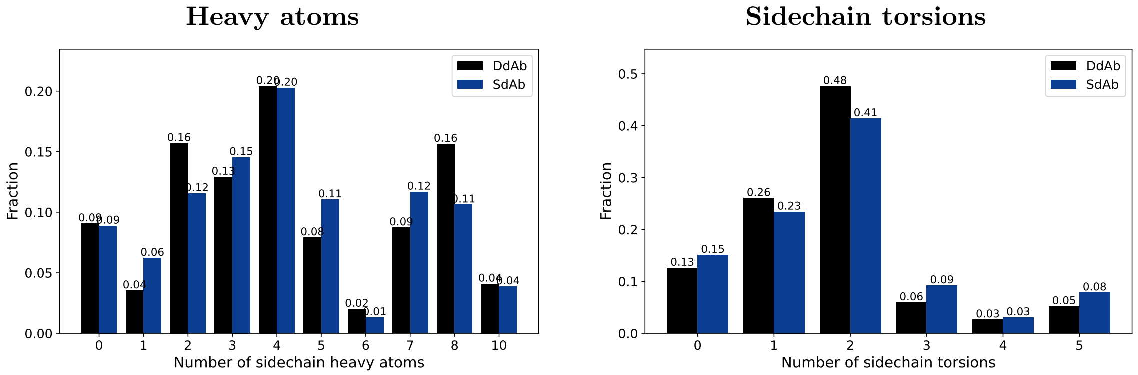


Figure S11: **Interface amino acids: distribution of side-chain sizes.** SdAb and DdAb display similar profiles. Dataset Ab-ALL, solvated model

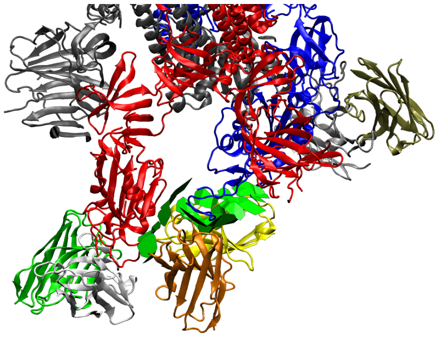

**(A)** pdbid: 7JV4

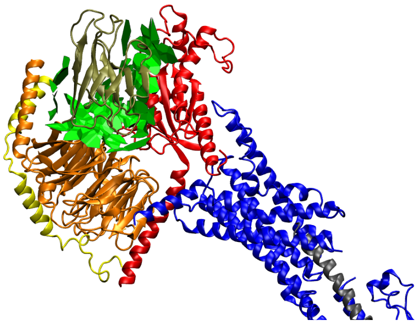

**B**: pdbid: 8jip

Figure S12: **Examples of highly fragmented antibody, target interfaces.** VMD screenshots of two antibody:target complexes displaying a large number of disconnected interface patches. These examples illustrate cases where the binding interface is spatially fragmented rather than forming a compact continuous surface.

### S2 SI: results

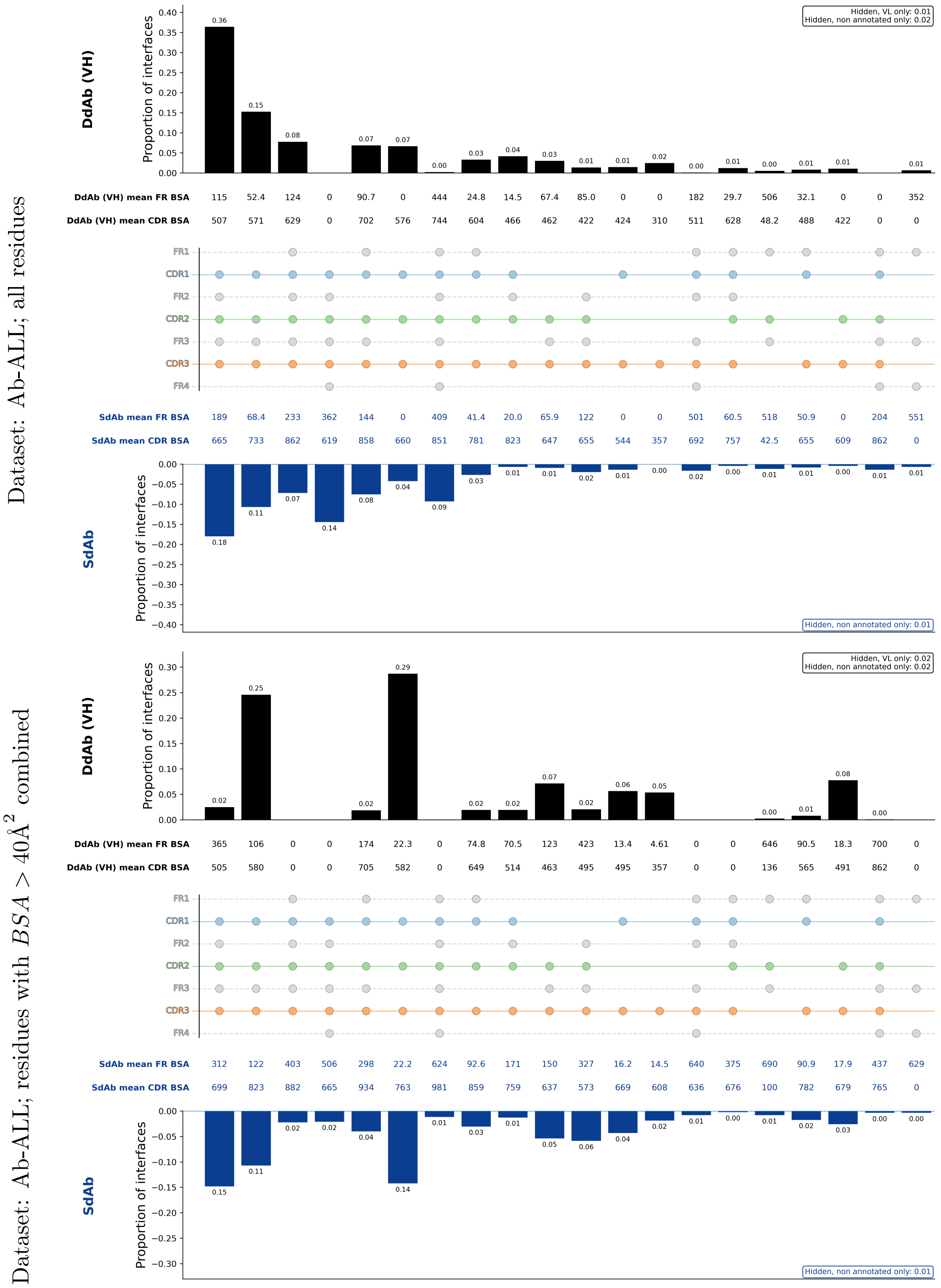


Figure S13: **Contact profiles. DdAb VH vs SdAb (Top)** Our solvated model–any residue contributing an interface atom is at the interface. **(Bottom)** Our model with a BSA threshold: residues within a region must contribute $\geq 40\text{Å}^2$ to be considered at the interface. Using a $40\text{Å}^2$ BSA threshold to define regional participation reduces the influence of *marginal* contacts. This yields profiles broadly consistent with the Cambridge analysis [14], although the agreement is not exact, with the main remaining differences observed for SdAb interfaces.

Dataset: Ab-ALL

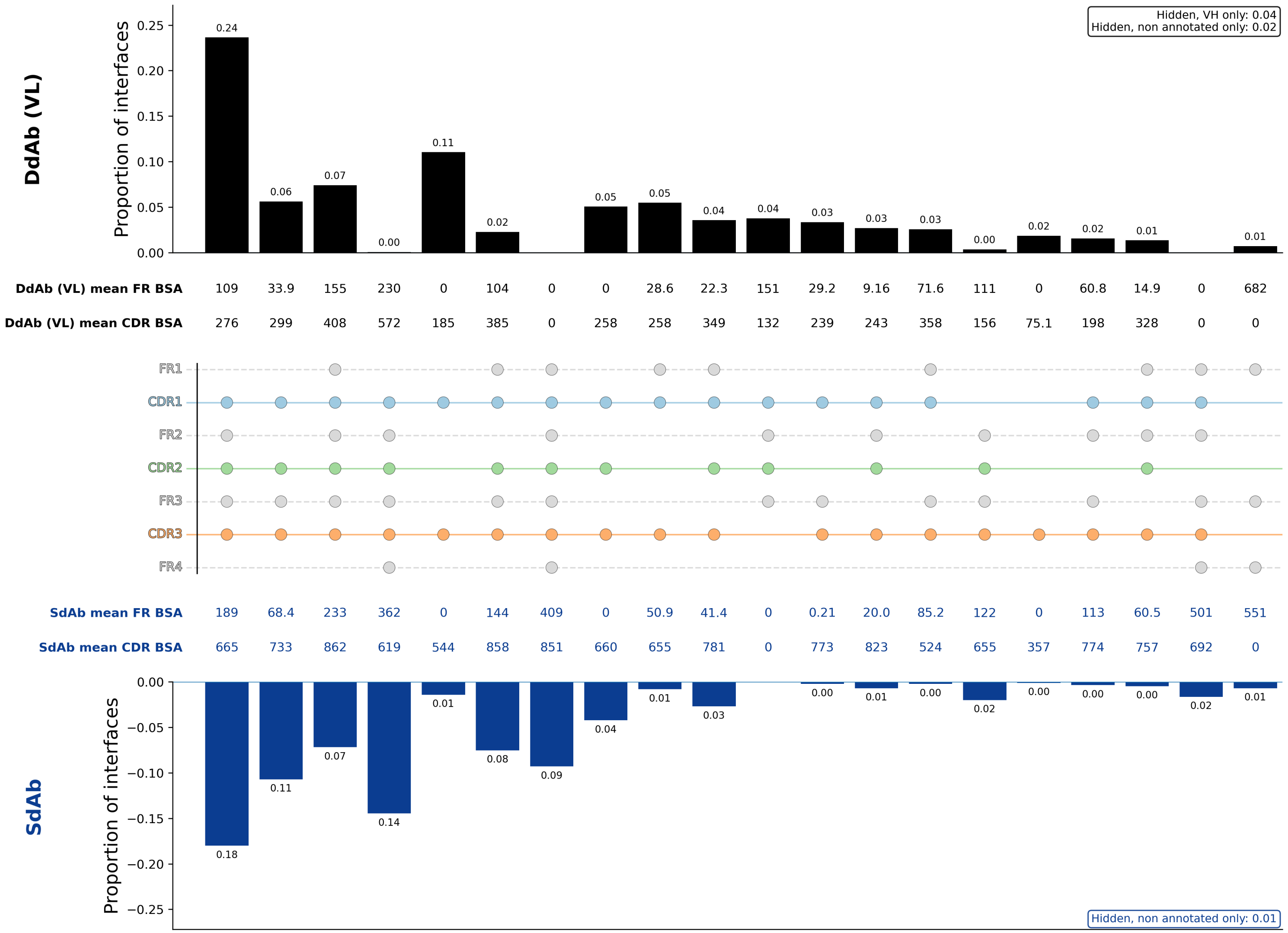


Figure S14: **Contact profiles DdAb VL vs SdAb for the dataset Ab-ALL.** A profile characterizes which regions of the light chain, out of CDRs and FRs, participate in the interface. The top 20 profiles (SdAb + DdAb) are displayed. The model used is solvated
